# Latent mutations in the ancestries of alleles under selection

**DOI:** 10.1101/2023.06.13.544872

**Authors:** Wai-Tong (Louis) Fan, John Wakeley

## Abstract

We consider a single genetic locus with two alleles *A*_1_ and *A*_2_ in a large haploid population. The locus is subject to selection and two-way, or recurrent, mutation. Assuming the allele frequencies follow a Wright-Fisher diffusion and have reached stationarity, we describe the asymptotic behaviors of the conditional gene genealogy and the latent mutations of a sample with known allele counts, when the count *n*_1_ of allele *A*_1_ is fixed, and when either or both the sample size *n* and the selection strength |*α*| tend to infinity. Our study extends previous work under neutrality to the case of non-neutral rare alleles, asserting that when selection is not too strong relative to the sample size, even if it is strongly positive or strongly negative in the usual sense (*α* → −∞ or *α* → +∞), the number of latent mutations of the *n*_1_ copies of allele *A*_1_ follows the same distribution as the number of alleles in the Ewens sampling formula. On the other hand, very strong positive selection relative to the sample size leads to neutral gene genealogies with a single ancient latent mutation. We also demonstrate robustness of our asymptotic results against changing population sizes, when one of |*α*| or *n* is large.

## 1. Introduction

The observed copies of a particular allele in a sample descend from an unknown number of distinct mutations. If *k*_1_ is the number of these ‘latent’ mutations for allele *A*_1_ when it is observed *n*_1_ times in a sample, then *k*_1_ ∈ {1, 2, …, *n*_1_}. Although latent mutations are not observed directly, they can be modeled as outcomes of the stochastic ancestral process of a sample and inferred from patterns of variation in DNA data (Harpak et al., 2016; Seplyarskiy et al., 2021; Johnson et al., 2022). Analytical results on the distribution and timing of latent mutations of rare neutral alleles are given in Wakeley et al. (2023). Here we consider non-neutral alleles which may be under strong selection and which may or may not be rare. We take two different approaches to modeling latent mutations under selection and recurrent mutation. The first approach uses the idea of coalescence in a random background of allele frequencies in the population (Barton et al., 2004). The second uses the conditional ancestral selection graph (Slade, 2000a) and demonstrates results consistent with those from the first approach.

Wakeley et al. (2023) also contains an application to the frequencies of single-nucleotide sites with counts *n*_1_ ∈ {1, 2, …, 40} of synonymous mutations in a subsample of 57K non-Finnish European individuals (*n* = 114K) from the *gnomAD* database (Karczewski et al., 2020). Dramatic differences in sample frequency distributions of rare alleles with different mutation rates, categorized by the ‘Roulette’ method of Seplyarskiy et al. (2023), were well explained by an empirical demographic model with recurrent mutation but no selection. Seplyarskiy et al. (2023, Fig. 3a) showed using simulations that a neutral, parametric demographic model fitted to these data also explained the frequencies of mutation in counts *n*_1_ ≤ 10^4^. Polymorphic sites with small mutation counts comprise the bulk of variation in humans. They represent a rich source of information about demographic history and possibly selection. Sites with *n*_1_ ∈ {1, 2, …, 40} make up about 95% of all polymorphic sites in the *gnomAD* data used in Wakeley et al. (2023).

At present humans are the only species with sufficient genomic data to apply such models of rare variants which rely on limiting approximations for large sample sizes. Whereas the neutral models in Wakeley et al. (2023) and Seplyarskiy et al. (2023) also account for the extreme population growth of humans (Keinan and Clark, 2012; Gazave et al., 2014; Gao and Keinan, 2016), in considering selection here we focus on populations of constant size. Previous theoretical work on populations of constant size has shown that distributions of rare alleles are in fact unaffected even by moderately strong selection (Joyce and Tavaré, 1995; Joyce, 1995). Specifically, the counts of latent mutations obey the independent Poisson statistics of rare alleles in the Ewens sampling formula (Ewens, 1972; Arratia et al., 1992, 2003). This is also the case in Wakeley et al. (2023) when the population size is constant. In the present work we investigate the robustness of these results to very strong selection. Theory also predicts that rare alleles tend to be young (Kimura and Ohta, 1973; Watterson, 1976). Mathieson and McVean (2014) and Platt et al. (2019) have demonstrated empirically that rare non-synonymous or otherwise functional alleles in the human genome are even younger than non-functional rare alleles. In the present work we also investigate how strong selection and rarity affect the ages of latent mutations.

We assume there are two possible alleles, *A*_1_ and *A*_2_, at a single genetic locus in a large haploid population. We begin by assuming that the population size *N* is constant over time. In Section 3.4 we consider time-varying population size. One allele or the other is favored by directional selection. Mutation is recurrent and happens in both directions. In the diffusion approximation, time is measured in proportion to *N*_*e*_ generations where *N*_*e*_ is the effective population size (Ewens, 2004). Under the Wright-Fisher model of reproduction, *N*_*e*_ = *N*. Under the Moran model of reproduction (Moran, 1958, 1962), *N*_*e*_ = *N/*2. With these assumptions, the frequency of *A*_1_ alleles is well approximated by a process *X* that solves (1) below and has parameters *θ*_1_, *θ*_2_ and *α* as *N* → ∞. For a haploid population, *θ*_*i*_ = 2*N*_*e*_*u*_*i*_ and *α* = 2*N*_*e*_*s*, in which *u*_*i*_ is the per-generation rate of *A*_3*−i*_ → *A*_*i*_ mutations and *s* is the selection coefficient. If there is no dominance, these results can be extended to diploids, in which case *θ*_*i*_ = 4*N*_*e*_*u*_*i*_ and *α* = 4*N*_*e*_*s*.

Thus, we assume that allele-frequency dynamics in the population obey the Wright-Fisher diffusion (Fisher, 1930; Wright, 1931; Ewens, 2004) with parameters *θ*_1_ and *θ*_2_ for mutations *A*_2_ → *A*_1_ and *A*_1_ → *A*_2_, respectively, and *α* for the selective advantage (if *α >* 0) or disadvantage (if *α <* 0) of allele *A*_1_. That is, we let *X*(*t*) be the relative frequency of *A*_1_ in the population at time *t*, and assume that its forward-time dynamics is described by the stochastic differential equation

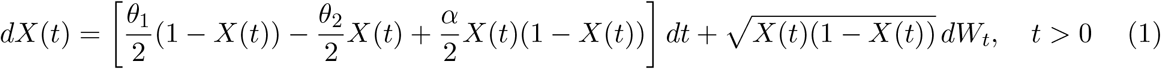

in which *W*_*t*_ is the Wiener process, also called the standard Brownian motion.

Both of the approaches (random background and ancestral selection graph) we take to modeling latent mutations rely on the assumption that the population has reached equilibrium, which occurs in the limit *t* → ∞. The stationary probability density of *X* is

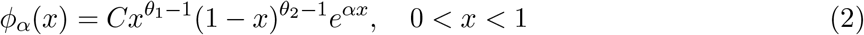

(Wright, 1931; Ewens, 2004). We explicitly denote the dependence on *α* because this parameter plays a key role in what follows. The normalizing constant *C* guarantees that 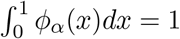. It is given by

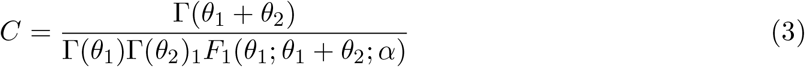

in which Γ(*a*) is the gamma function and _1_*F*_1_(*a*; *b*; *z*) is the confluent hypergeometric function, or Kummer’s function; see Abramowitz and Stegun (1964) and Slater (1960).

By definition, latent mutations occur in the ancestry of a sample. When a sample of total size *n* is taken from a population with stationary density (2), it will contain a random number 𝒩_1_ of copies of allele *A*_1_ and 𝒩_2_ = *n* − 𝒩_1_ copies of allele *A*_2_. The probability that 𝒩_1_ is equal to *n*_1_ is equal to

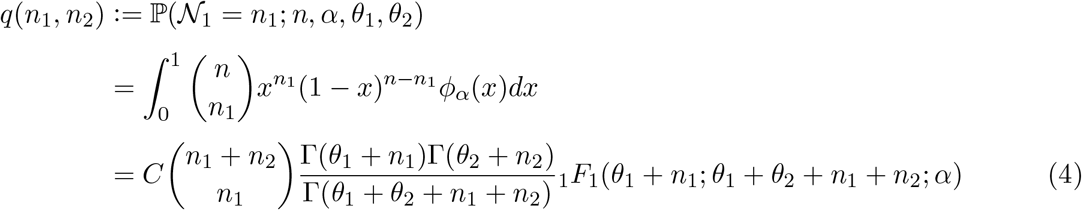

for *n*_1_ ∈ {0, 1, …, *n*} and *n*_2_ = *n* − *n*_1_, and with *C* again given by (3). The notation *q*(·) is from Slade (2000a,b) and is convenient for the ancestral selection graph.

Suppose now we are given the sample count, that is, we know that among the *n* uniformly sampled haploid individuals, *n*_1_ of them are of type 1 and the remaining *n*_2_ = *n* − *n*_1_ are of type 2. Then the posterior density of the population frequency of *A*_1_ conditional on the sample is

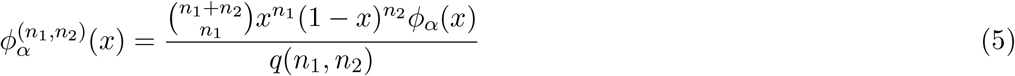

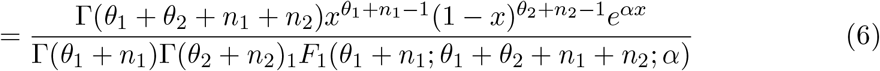

from Bayes’ theorem with prior density *ϕ*_*α*_.

The sampling probability *q*(*n*_1_, *n*_2_) in (4) and the resulting posterior density 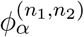 play major roles in the two approaches we take to modeling latent mutations. Specifically, transition probabilities in the conditional ancestral selection graph depend on ratios of sampling probabilities (Slade, 2000b) and the allele frequency in the ancestral process of Barton et al. (2004) has initial density 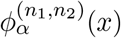 when conditioned on the sample.

We describe the occurrence of latent mutations in the ancestry of allele *A*_1_ conditional on the sample count *n*_1_. We say that *A*_1_ is *rare* when the sample size *n* is much larger than *n*_1_. We enforce this rarity of *A*_1_ by letting *n*_2_ ∼ *n* tend to infinity with *n*_1_ fixed, or finite. We present some results for cases in which *A*_1_ is not rare in this sense, that is when neither *n*_1_ nor *n*_2_ is large. In this case we also describe the conditional ancestry of *A*_2_, but overall our focus is on large samples and rare *A*_1_. This is the same, sample-based concept of rarity that was used in Wakeley et al. (2023) and previously considered by Joyce and Tavaré (1995) and Joyce (1995). It may be distinguished from rarity in the population, though of course finding *A*_1_ rare in a large sample is most likely when the population frequency *x* is small.

By *strong selection* we mean large |*α*|. We model rarity and strong selection together under the assumption that 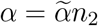 for some constant 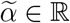. We study latent mutations and the ancestral processes which generate them under three scenarios: (i) |*α*| large with *n*_2_ fixed, (ii) *n*_2_ large with *α* fixed, and (iii) both |*α*| and *n*_2_ large with *α* = *α/n*_2_ fixed. In making approximations for large *n*_2_ and/or large |*α*|, we make extensive use of asymptotic results for ratios of gamma functions and for the confluent hypergeometric function which are presented in Appendix A.

The parameters *θ*_1_ and *θ*_2_ are fixed constants throughout, with *θ*_1_, *θ*_2_ *>* 0. For single nucleotide sites, these population-scaled mutation rates have been estimated for many species, using average pairwise sequence differences and assuming constant population size, and are typically about 0.01 with a range of about 0.0001 to 0.1 (Leffler et al., 2012). Values for humans are smaller but they vary almost as widely among sites in the genome, with a mean of about 0.0008 and a range of about 0.0001 to 0.02 (Seplyarskiy et al., 2021, 2023; Wakeley et al., 2023). In contrast, there is no reason to suppose that the selection parameter |*α*| is small (Eyre-Walker and Keightley, 2007; Chen et al., 2020; Agarwal et al., 2023). Note that our introduction of a constant 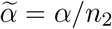 is simply a device to specify the relative importance of rarity as opposed strong selection, not a hypothesis about biology.

The case of a rare neutral allele was considered in Wakeley et al. (2023) where it was shown that the number of latent mutations in the ancestry of the *n*_1_ copies of allele *A*_1_ follows the same distribution as the number of alleles in the Ewens sampling formula (Ewens, 1972) with sample size *n*_1_ and mutation parameter *θ*_1_. Let *K*_1_ be the random number of these latent mutations for allele *A*_1_ in the ancestry of the sample. Further, let *ξ*_*j*_ be a Bernoulli random variable with probability of success

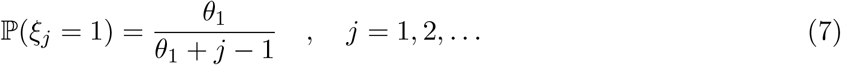

Under neutrality for large sample size and conditional on 𝒩_1_ = *n*_1_,

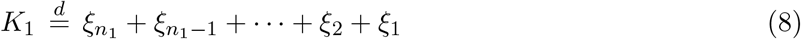

which gives the stated Ewens sampling result (Arratia et al., 1992). In (8) and below, 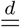 denotes equal in distribution. Note that, because coalescence is among exchangeable lineages, the full Ewens sampling formula should apply if we were to keep track of the sizes of latent mutations; see Crane (2016) and Tavaré (2021) for recent reviews.

Here we apply the model of coalescence in a random background described by Barton et al. (2004) to prove these results (7) and (8) for rare alleles in large samples and especially to extend the analysis of latent mutations to scenarios which include selection. We investigate both the number of latent mutations and their timing in the ancestry of the sample, and we allow that selection may be strong. We also show how the same scenarios can be treated using the conditional ancestral selection graph (Slade, 2000a), giving the same limiting results for all three scenarios.

Briefly, we find that positive selection does not in general lead to (7) and (8), that very strong positive selection (relative to the sample size) leads to neutral gene genealogies with a single ancient latent mutation for the favored allele. This is described in Section 3 for scenario (i) and for the case 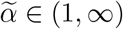 in scenario (iii). On other hand, when selection is not too strong relative to the sample size, then extreme rarity of *A*_1_ in the sample can effectively override strong positive selection and retrieve (7) and (8). This is described in Section 3 for scenario (ii) and for the case 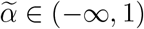 in scenario (iii). Figures 1, 2 and 3 illustrate our results in the three scenarios.

**Figure 1:**
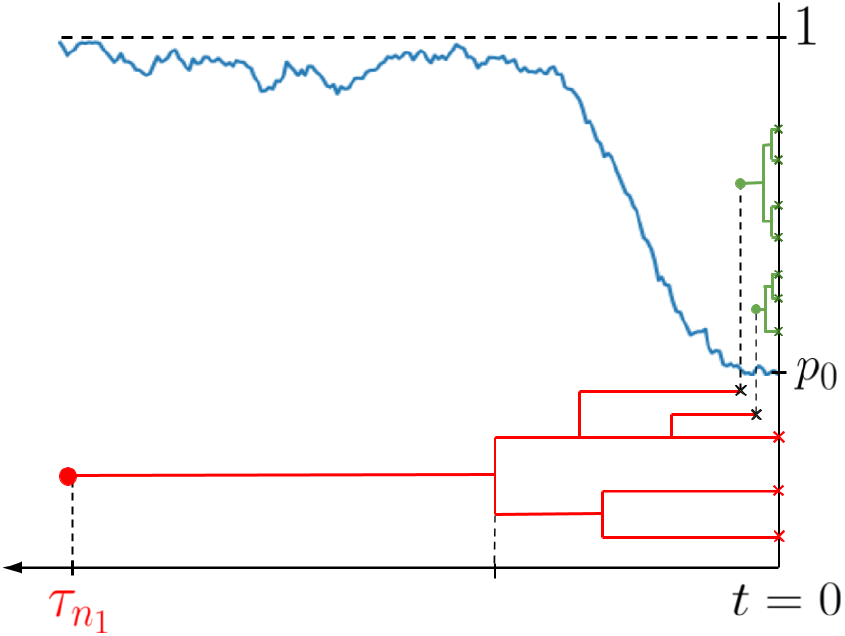
Conditional genealogy of a sample with (*n*_1_, *n*_2_) = (3, 7) at the present time *t* = 0. The fluctuating blue curve shows the process (*p*_*t*_)*t∈*ℝ_+_ of the population frequency of type 1 backward in time. In this example, *p*_*t*_ approaches 1 from *p*_0_ and the 7 type-2 lineages coalesce and mutate, producing an additional *K*_2_ = 2 type-1 lineages. The 5 = 3 + 2 type-1 lineages then coalesce without mutating, reaching their common ancestral lineage at time 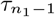 and finally mutating at time 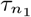. Under scenario (i), that is when *α* is large: *p*_0_ will already be close to 1, coalescence and mutation among the type-2 lineages will occur quickly, with *K*_2_ according to the Ewens sampling formula, coalescence among the type-1 lineages will follow the Kingman coalescent, and 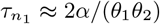.

**Figure 2:**
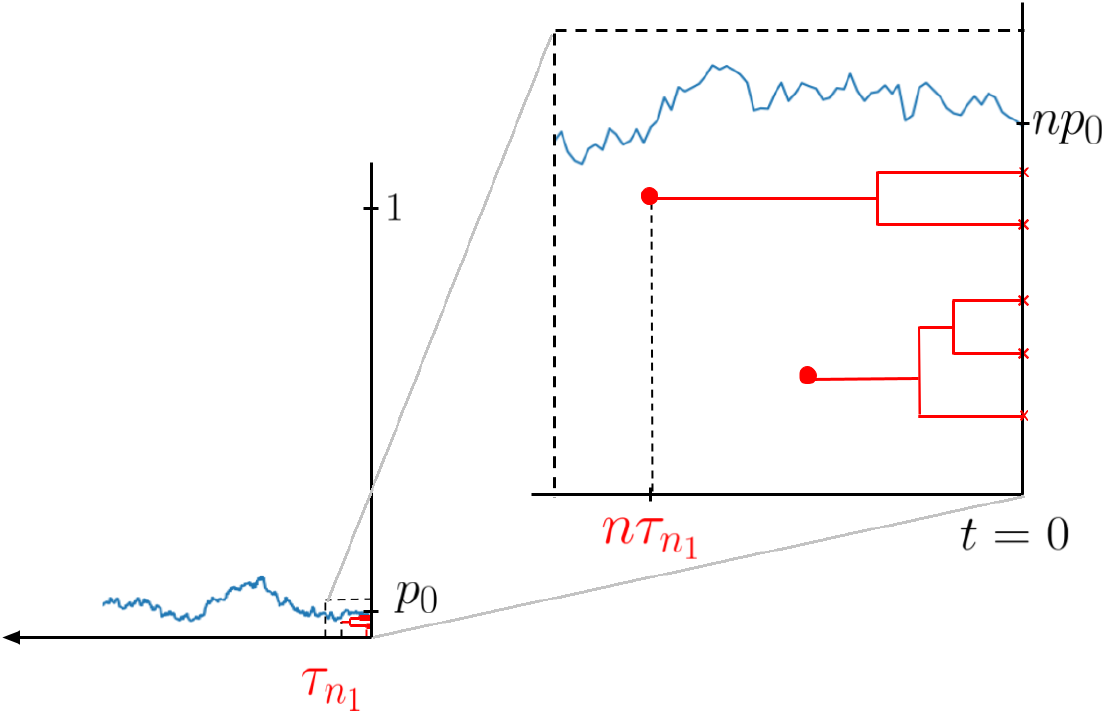
Conditional genealogy of a sample with observed frequencies (*n*_1_, *n*_2_) at the present time *t* = 0, where *n*_1_ = 5 and *n*_2_ is large, and 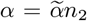 for a constant 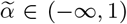. The *n*_2_ samples are not shown. In this figure, *K*_1_ = 2 and the two red bullets are mutation events from type 1 to type 2. In scenario (iii), *K*_1_ is distributed like the number of alleles in the Ewens sampling formula, and the timing of the type-1 events are small (of order *O*(1*/n*) on the coalescent time scale). The rescaled process 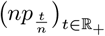 is well approximated by the diffusion process (40) with initial distribution 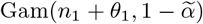.

**Figure 3:**
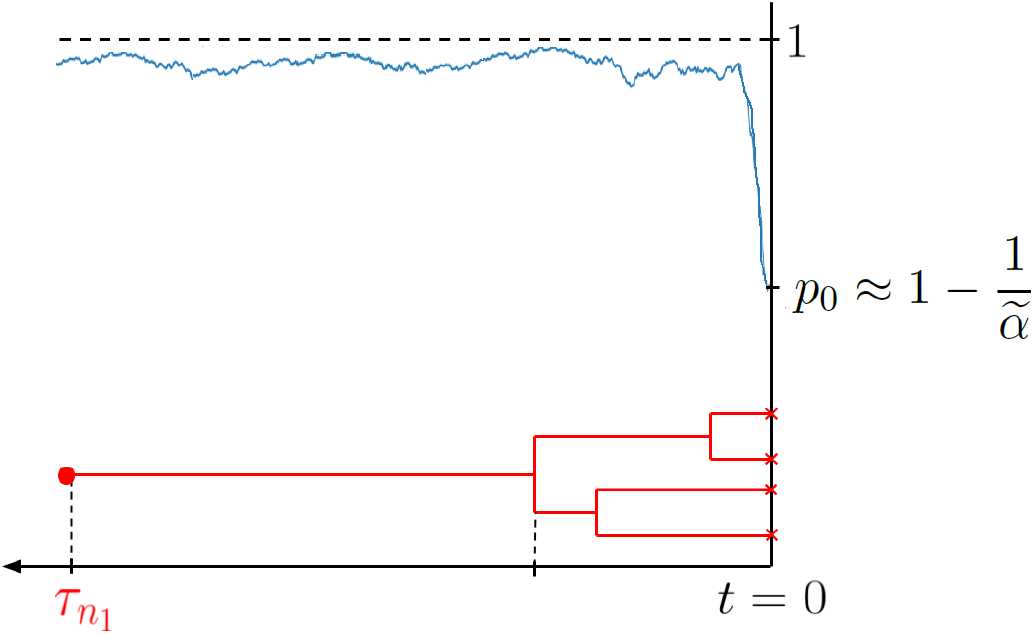
Conditional genealogy of a sample with observed frequencies (*n*_1_, *n*_2_) at the present time *t* = 0, where *n*_1_ = 4 and *n*_2_ is large, and 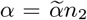 for a constant 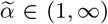. The *n*_2_ samples are not shown. This scenario is reminiscent of scenario (i) if we focus on the genealogy of only the *A*_1_ lineages. These lineages first coalesce as the Kingman coalescent (without mutation) until there is only one lineage, which take *O*(1) amount of time. Then it takes *O*(*n*_2_) amount of time for the single latent mutation to occur at time 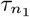.

We note that Favero and Jenkins (2024) have recently performed detailed analysis of a *d*-allele diffusion model, where the selective advantage of one allele grows to infinity and the other parameters remain fixed. Their findings confirm and extend what we establish for scenario (i) in the two-allele model in Sections 3.1 and 4.1. In addition, Favero and Jenkins (2024) prove the duality of the strong-selection limit of the diffusion and the corresponding ancestral selection graph.

## 2. Sample frequencies and posterior population frequencies

In this section, we present asymptotic results for the sampling probability *q*(*n*_1_, *n*_2_) in (4) and the posterior density 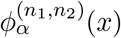 in (5) in our three regimes of interest: (i) |*α*| large with *n*_2_ (ii) *n*_2_ large with *α* fixed, and (iii) both |*α*| and *n*_2_ large with 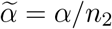 fixed.

### 2.1 Asymptotics for sampling probabilities

In the case of strong selection and moderate sample size, that is |*α*| large with *n*_2_ fixed, applying (A.4a) and (A.4b) to (4) gives

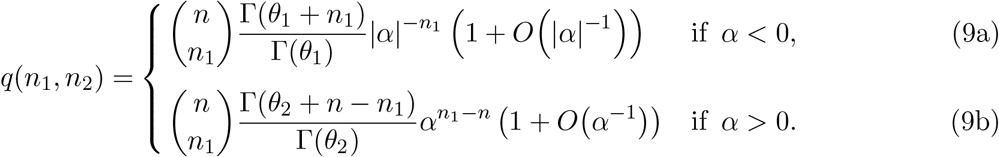

Here we focus on the leading-order terms but note that the next-order terms are straightforward to obtain using (A.4a) and (A.4b) and additional higher-order terms could be computed using (4.1.2) and (4.1.6) in Slater (1960). In (9a), each additional copy of *A*_1_ decreases the sampling probability by a factor of 1*/*|*α*| so the most likely sample is one which contains no copies of *A*_1_. In (9b), each additional copy of *A*_1_ increases the sampling probability by a factor of *α* so the most likely sample is monomorphic for *A*_1_. However, these results are perfectly symmetric for the two alleles. Switching allelic labels and swapping |*α*| for *α* changes (9a) into (9b). That is, allele *A*_2_ experiences the same effects of positive/negative selection in (9a)/(9b) as the focal allele *A*_1_ does in (9b)/(9a).

In the case of large sample size and moderate selection, that is *n*_2_ large with *α* fixed, applying (A.5) to (4) gives

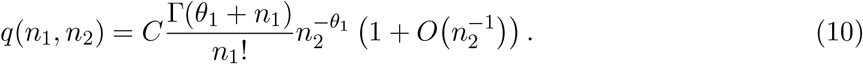

This has the same form as the neutral result, equation (22) in Wakeley et al. (2023), only with the additional factor _1_*F*_1_(*θ*_1_; *θ*_1_ + *θ*_2_; *α*) in the denominator of the constant *C*. With respect to the count of the focal allele *A*_1_, the distribution is similar to a (degenerate) negative-binomial distribution with parameters *p* = 1*/n*_2_ and *r* = *θ*_1_, like the corresponding result in Theorem 2 of Watterson (1974) for neutral alleles which propagate by a linear birth-death process. The effect of selection is only to uniformly raise or lower the chances of seeing *n*_1_ copies of *A*_1_ in a very large sample. The additional factor _1_*F*_1_(*θ*_1_; *θ*_1_ + *θ*_2_; *α*) in the denominator of *C* is a decreasing function of *α*, which is equal to 1 when *α* = 0 and approaches 0 quickly from there as *α* increases. Greater selection against (respectively, for) *A*_1_ increases (respectively, decreases) the chance of it being rare but does not affect the shape of the distribution of *n*_1_, at least to leading order in 1*/n*_2_.

In the case of large sample size and strong selection, that is both |*α*| and *n*_2_ large with 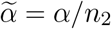 fixed, applying (A.4a), (A.4b), (A.6a), (A.6b) to (4) gives

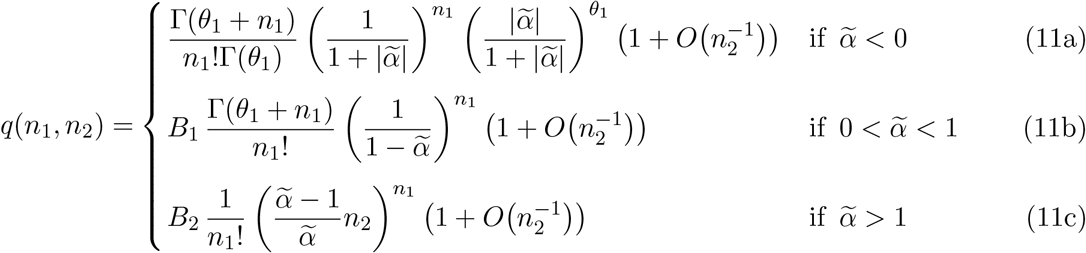

with constants *B*_1_ and *B*_2_ which are unremarkable except in their dependence on *n*_2_:

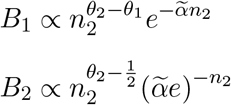

such that *q*(*n*_1_, *n*_2_) becomes tiny as *n*_2_ grows. In (11b) and (11c), allele *A*_1_ is favored by selection so it will be unlikely for its sample count to be very small.

To see how (11b) and (11c) compare to (11a), consider how these three sampling probabilities change as *n*_1_ increases:

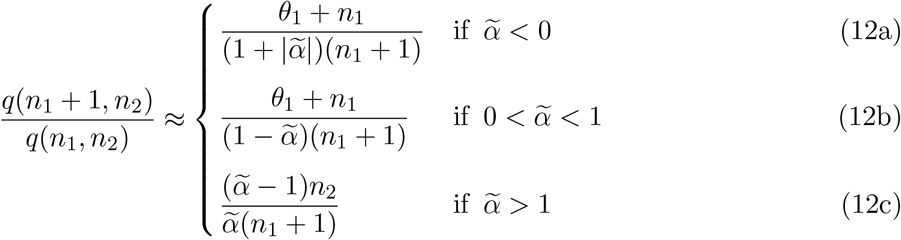

where the approximation is for large *n*_2_, i.e. omitting the 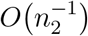 parts of (11a), (11b) and (11c). The first two differ from the corresponding neutral result (*θ*_1_ + *n*_1_)*/*(*n*_1_ + 1) by the constant factors 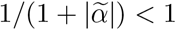 in (12a) and 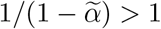 in (12b). Note that (10) gives the neutral result, as do (12a) and (12b) as 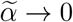. Relative to this, negative selection in (12a) makes additional copies of *A*_1_ less probable whereas positive selection in (12b) makes them more probable. But (12a) and (12b) differ from the neutral result only by these constant factors. Equation (12c) is quite different. With 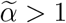, each additional copy of *A*_1_ increases the sampling probability by a large factor, proportional to *n*_2_, making this case similar to the case of strong positive selection in (9b). This is as expected. What is surprising is (12b), namely that strong selection (*α* → ∞) in favor of *A*_1_ can be made to resemble neutrality simply by increasing the sample size relative to *n*_1_.

#### 2.1.1. Comparison to discrete Moran and Wright-Fisher models

We emphasize that our analyses in this work are of the Wright-Fisher diffusion model, given here as the SDE (1) with stationary density (2). It is of interest to know how well our results hold for discrete, exact models such as the Moran model and the Wright-Fisher model, especially as |*α*| → ∞ or *n* → ∞ for finite *n*_1_, in which cases we might expect the diffusion to be a relatively poor description of the dynamics. In this section, we focus on (11a) and show that it can be obtained in a different way from a discrete-time Moran model, without first passing to the diffusion limit, but that this cannot be done in general starting from the discrete-time Wright-Fisher model.

To leading order in 1*/n*_2_, (11a) is identical to the probability mass function of a negative binomial distribution with parameters 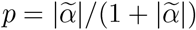 and *r* = *θ*_1_. Charlesworth and Hill (2019) found this same result starting from the strong-selection approximation which Nei (1968) had obtained for the diffusion model of Wright (1937). Here selection *against A*_1_ is so strong that it never reaches appreciable frequency in the population. In the limit, or ignoring the 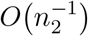 part in (11a), this distribution sums to one over all *n*_1_ ∈ {0, 1, 2, …}. The corresponding sum for the degenerate distribution in (10) diverges, because under neutrality there is a non-trivial chance that *A*_1_ reaches appreciable frequency in the population.

Consider a discrete-time haploid Moran model with population size *N*, in which allele *A*_2_ is favored by selection. Specifically, *A*_1_ and *A*_2_ have equal chances of being chosen to reproduce but different chances of being chosen to die: each *A*_1_ has an increased chance 1 + *s* compared to each *A*_2_. Upon reproduction, the offspring of an *A*_*i*_, *i* ∈ {1, 2}, has type *A*_*i*_ with probability 1 − *u*_3*−i*_ and the other type *A*_3*−i*_ with probability *u*_3*−i*_. If there are currently *ℓ* copies of *A*_1_ and *N* − *ℓ* copies of *A*_2_, then in the next time step there will *ℓ* + 1 copies of *A*_1_ with probability

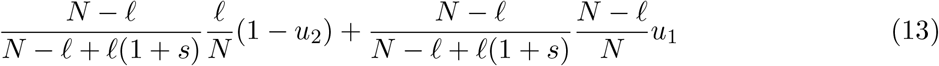

and *ℓ* − 1 copies of *A*_1_ with probability

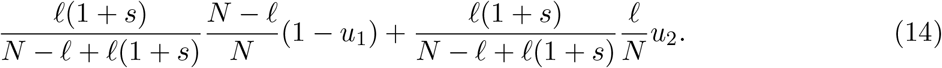

The fraction of *A*_1_ converges to the Wright-Fisher diffusion process (1) as *N* → ∞ if time is measured in units of *N* (*N* − 1)*/*2 discrete steps, i.e. *dt* = 2*/N* (*N* − 1), with *u*_1_ = *θ*_1_*/N, u*_2_ = *θ*_2_*/N* and *s* = −*α/N*.

As another way of obtaining (11a), we assume that *s* ≫ *u*_1_, *u*_2_. In particular, let *Nu*_1_ → *θ*_1_ and *Nu*_2_ → *θ*_2_ as *N* → ∞ just as in the diffusion model, but let *s* be a constant. Then we may appeal to the analogous model and limit process (iii) of Karlin and McGregor (1964) which had no selection but instead assumed that *u*_2_ ≫ *u*_1_. Similarly here we expect that allele *A*_1_ will be held in negligible relative frequency in the population and instead be present in a finite number of copies as *N* → ∞, only here due to strong selection rather than strong mutation.

In view of this scaling of the mutation rates by *N* and for comparison with the Wright-Fisher model below, we rescale time so that it is measured in unit of generations, or *N* time steps. Then with *dt* = 1*/N*, we can rewrite (13) and (14) as

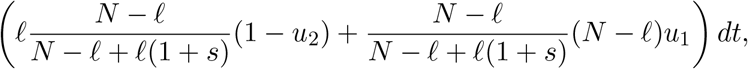

and

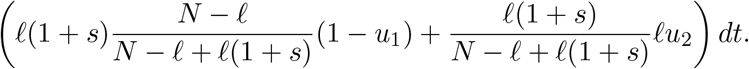

Then in the limit *N* → ∞, (13) and (14) describe to a continuous-time process in which

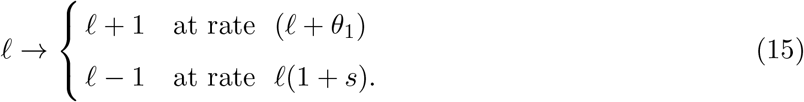

In other words, the number of copies of *A*_1_ in the population evolves according to a birth-death process with immigration where the birth rate is *λ* = 1, the death rate is *μ* = 1 + *s* and the immigration rate is *κ* = *θ*_1_. From (52) in Kendall (1949), the distribution of the number of copies of *A*_1_ in the population at stationarity will be negative binomial with parameters 1 − *λ/μ* = *s/*(1 + *s*) and *κ/λ* = *θ*_1_, or

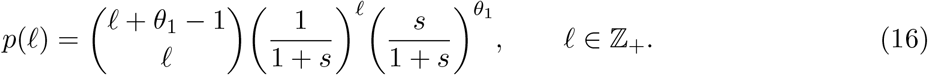

In getting to (11a) above, which we note is for *α <* 0, we first applied the diffusion limit then let the selection parameter |*α*| be large, specifically proportional to the number *n*_2_ of copies of *A*_2_ in a sample of large size *n* for a given fixed number *n*_1_ of copies of *A*_1_. In the current haploid Moran model with selection against *A*_1_, |*α*| = *Ns*, and the scalar 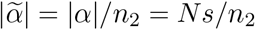. Define *a* := *n*_2_*/N*. Then 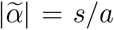 and we may think of *a* as (close to) the proportion of the population sampled, because *n*_2_*/N* ∼ *n/N*.

If the number of copies of *A*_1_ in the population is *ℓ*, then the probability there are *n*_1_ copies in a sample of size *n* taken without replacement from the total population of size *N* is given by the hypergeometric distribution

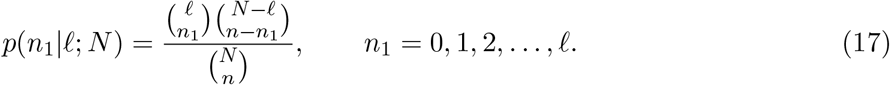

Since *n* − *n*_1_ = *n*_2_ = *aN* and taking *N* → ∞, (17) converges to the binomial distribution

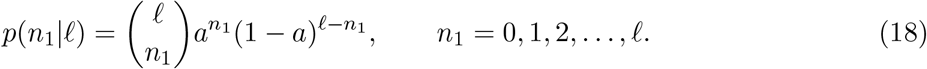

This gives another route to (11a), namely using (16) and (18), and setting *y* = *ℓ* − *n*_1_,

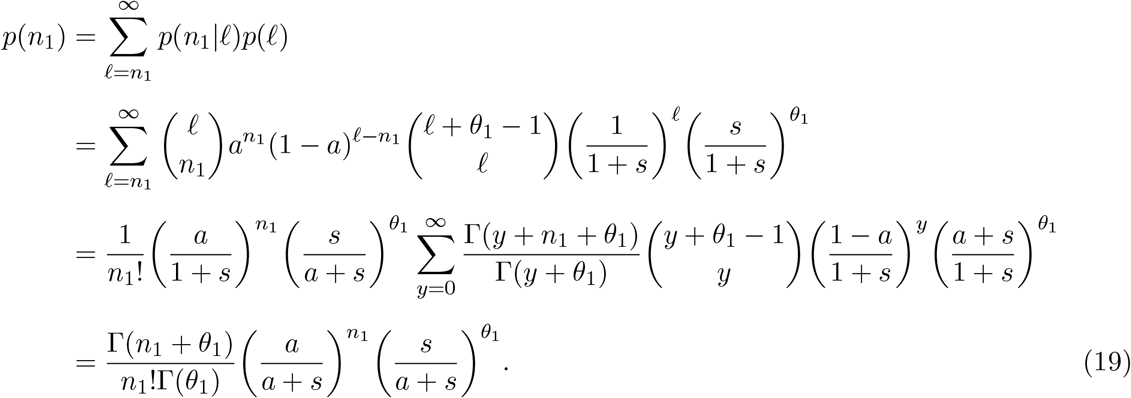

The end result (19) is equal to the leading order part of (11a) since 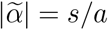.

We can contrast this with a discrete-time haploid Wright-Fisher model with population size *N*, in which time is already measured in generations. Under the same assumptions that gave (15), namely *s* constant and *u*_1_, *u*_2_ ∝ 1*/N* as *N* → ∞, we can use equation (33) in Nagylaki (1990) which specifies that, conditional on the number *ℓ*_*g*_ of copies of *A*_1_ in generation *g*, the number *ℓ*_*g*+1_ has the Poisson distribution

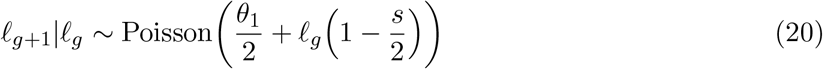

with *θ*_1_ := 2*Nu*_1_. Now *ℓ* evolves by a Poisson branching process with Poisson immigration rather than by the birth-death process with immigration in (15). Although here too the number of copies of *A*_1_ in the population will converge to a stationary distribution (Heathcote, 1965), it will not in general be a negative binomial distribution. Thus (11a) is consistent with the per-generation dynamics of rare alleles in the Moran model but not in the Wright-Fisher model.

### 2.2 Asymptotics for population frequencies conditional on the sample

Next, we obtain asymptotics for the posterior probability density 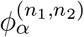 in (6). Let 𝒫 ([0, 1]) be the space of probability measures on [0, 1] endowed with the weak convergence topology (i.e with test functions in the space *C*_*b*_([0, 1]) of bounded continuous functions on [0, 1]).

#### Lemma 1.

*Let n*_1_ ∈ ℕ *be fixed. The following convergences in 𝒫* ([0, 1]) *hold*.

i. *Suppose n* ∈ ℕ *is fixed and α* → ∞. *Then* 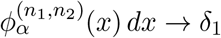.
ii. *Suppose α* ∈ ℕ *is fixed and n* → ∞. *Then* 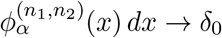.
iii. *Suppose* 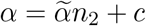 *where* 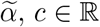 *are fixed and n*_2_ → ∞. *Then*

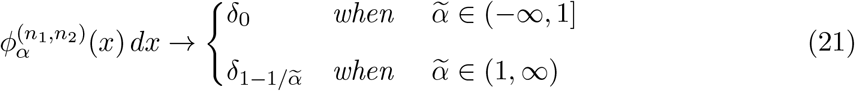

*where δ*_*x*_ *is the Dirac delta measure*.

The proof of Lemma 1 is given in Appendix B.

## 3. Conditional coalescence in a random background

In this section, we extend the approach of coalescence in a random background in Barton et al. (2004) to study the number and timing of latent mutations and other asymptotic properties of the conditional gene genealogy given the sample frequencies of *A*_1_ and *A*_2_. We also extend our results to time-varying populations in Section 3.4. While the setting of Barton et al. (2004) covers the case of a neutral locus linked to the selected locus, here we focus on the selected locus.

Suppose we are given a sample from the selected locus at the present time *t* = 0, and that we know the allelic types of the sample but we do not know how the sample was produced. What is the genealogy of the sample? This question was answered by Barton et al. (2004), who modeled the ancestral process using the structured coalescent with allelic types as subpopulations. The structured coalescent can be a model of subdivision with migration between local populations (Takahata, 1988; Notohara, 1990; Herbots, 1997) or a model of selection with mutation between allelic types (Kaplan et al., 1988; Darden et al., 1989). For samples from a population at stationarity as in Section 1, Barton et al. (2004) proved that this could be done rigorously starting with a Moran model with finite *N* then passing to the diffusion limit. Barton and Etheridge (2004) explored some properties of gene genealogies under this model, and Etheridge et al. (2006) used the same idea to describe genetic ancestries following a selective sweep.

Even if the sample frequencies are known, the allele frequencies in the population are unknown. A key feature of this method is to model allele-frequency trajectories backward in time. As pointed out by Barton et al. (2004), the Moran model with finite *N* is *reversible*, meaning that at stationarity the time-reversed process is the same (in distribution) as the forward-time Moran process. This is not a property of the Wright-Fisher model with finite *N* but does hold for their shared diffusion limit (1) with stationary density (2); see for instance Millet et al. (1989) for why this holds. Figure 1 gives an illustration of a genealogy with mutations and allele frequencies varying backward in time.

Looking backward in time, let 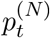 be the fraction of type 1 in the population and 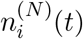 be the number of ancestral lineages of type *i* ∈ {1, 2} at time *t*. From Barton et al. (2004, Lemma 2.4), under the Moran model with stationary distribution, 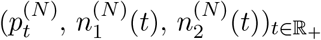 is a Markov process for each fixed *N*. Furthermore, Barton et al. (2004, Theorem 5.1) describes the joint convergence of the processes 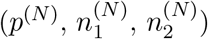 as *N* → ∞.

### Lemma 2

(Lemma 2.4 and Theorem 5.1 of Barton et al. (2004)). *Let* 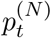 *and* 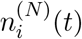 *be the fraction of type 1 in the population and the number of ancestral lineages of type i respectively, at time t backward, under the stationary Moran model. Then* 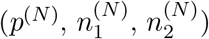 *is a Markov process for each N* ∈ ℕ. *As N* → ∞, *this process converges in distribution in the Skorohod space D*(ℝ_+_, [0, 1] × 𝕫_+_ × 𝕫_+_) *to a Markov process* (*p*_*t*_, *n*_1_(*t*), *n*_2_(*t*))_*t∈*ℝ+_ *described as follows:*

i. (*pt*) _*t∈*ℝ+_ *is a solution to equation* (1) *with stationary initial density ϕ*_*α*_. *In particular, it does not depend on* (*n*_1_(*t*), *n*_2_(*t*))_*t∈*ℝ+_.
ii. *Suppose the current state is* (*p, m*_1_, *m*_2_). *Then* (*n*_1_(*t*), *n*_2_(*t*))_*t∈*ℝ+_

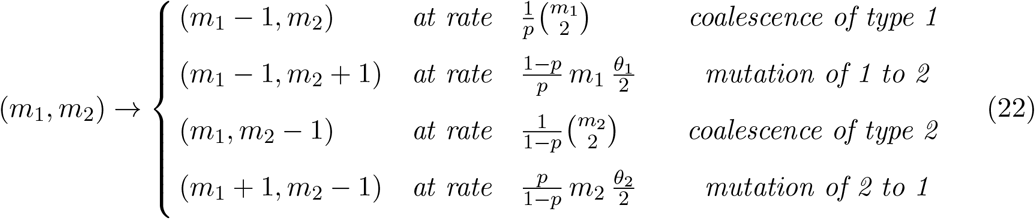

We compare the notation here with that of Barton et al. (2004) and Etheridge (2011). Type 1 here is their type P, type 2 here is their type Q. So, 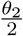 (resp. 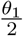) here is *μ*_1_ (resp. *μ*_2_) in Barton et al. (2004), and is *v*_1_ (resp. *v*_2_) in Etheridge (2011, eqn. (2.11)). The rescaled selection coefficient 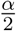 here is *s* in Barton et al. (2004). The Moran model in Barton et al. (2004) has population size 2*N* and each of the (2*N*)(2*N* − 1)*/*2 unordered pairs is picked to interact (one dies and immediately the other reproduces) at rate 1*/*2. Therefore, the diffusion limit in Barton et al. (2004, Lemma 3.1) is slower by a factor of 1*/*2 than the limit we consider here in this paper.

The proof of Barton et al. (2004, Theorem 5.1) leads to more information for the limiting process. Let 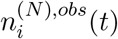 be the number of type *i* lineages at backward time *t* which are ancestral to the 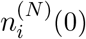 observed in the sample. Thus 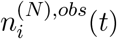 is non-increasing in *t*, but 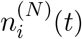 can increase as *t* increases due to mutations from type 3 − *i* to type *i*. Clearly, 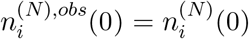. Note that a mutation from type 3 − *i* to type *i* backward in time corresponds to a mutation from type *i* to type 3 - *i* forward in time. To keep track of the number of mutation events versus the number of coalescent events, we let 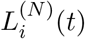 be the total number of latent mutations for type *i* during [0, *t*] backward in time. We have the following generalization of Lemma 1.

### Lemma 3

(Joint convergence). *Under the stationary Moran model, the backward process*

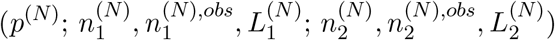

*is a Markov process for each fixed N* ∈ ℕ. *As N* → ∞, *this process converges in distribution in the Skorohod space* 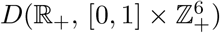 *to a continuous-time Markov process*

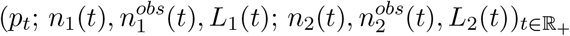

*such that*

i. (*p*_*t*_)_*t∈*ℝ+_ *is a solution to equation* (1) *with stationary initial density ϕ*_*α*_. *In particular, it does not depend on* 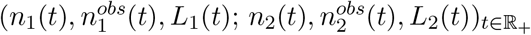.
ii. *At state* (*p*; *m*_1_, *a*_1_, *ℓ*_1_; *m*_2_, *a*_2_, *ℓ*_2_), *the process* 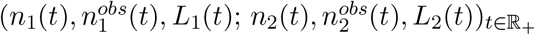 *evolves as* (*m*_1_, *a*_1_, *ℓ*_1_; *m*_2_, *a*_2_, *ℓ*_2_) →

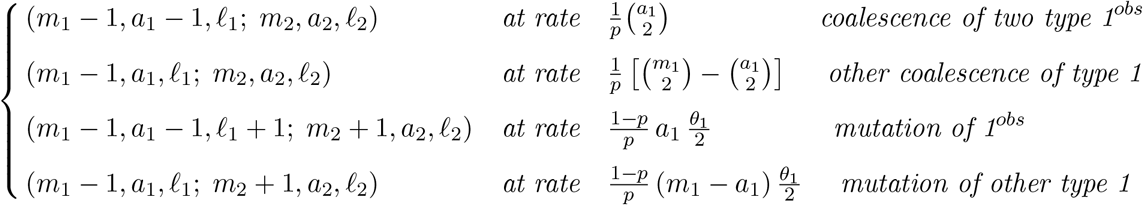

*and, similarly*,

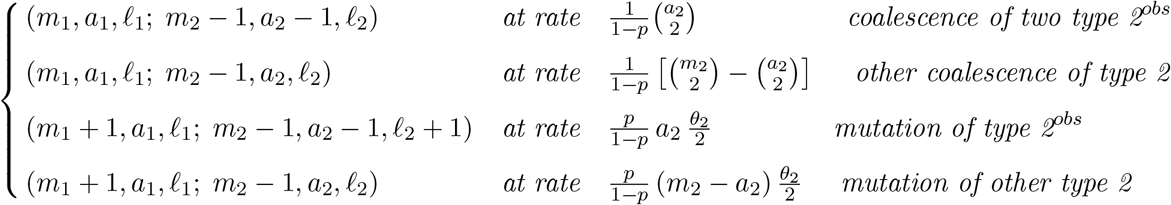

Now suppose that in addition to knowing the sample counts *n*_1_ and *n*_2_, we also know that these are the outcome of uniform random sampling, as in (4). Let ℙ_**n**_ be the conditional probability measure of the ancestral process in Lemma 1 including both *p*_*t*_ and the lineage dynamics, given that a uniformly picked sample has allelic counts **n** = (*n*_1_, *n*_2_). Under ℙ_**n**_, the limiting process in Lemma 1 has initial frequency 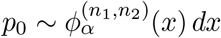 given by (6), and 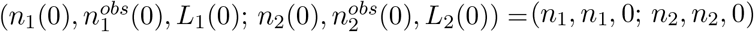. This follows from Bayes’ theorem, because by part (i) of Lemmas 2 and 3, the initial frequency has prior density given by (2).

Focus on type 1 for now. We care about the sequence of events (coalescence and mutation) backward in time for type 1, and the timing of these events. At each of these events, the number of type 1 lineages decreases by 1, either by coalescence or by mutation from type 1 to type 2. Hence 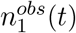 is non-increasing, but *n*_1_(*t*) can increase over time (backward) due to mutations from type 2 to type 1 (see Figure 1 for an illustration). Furthermore, the difference 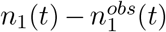 is the number of type 1 at time *t* that came from lineages that are of type 2 in the sample (at *t* = 0). We do not care about these 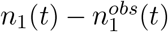 lineages, nor the mutation events from type 2 to type 1. Analogous considerations hold for type 2.

From Lemma 1, we immediately obtain the following simplified description for the conditional ancestral process in the limit *N* → ∞ for the two types. This description is the starting point of our analysis for constant population size; later in Proposition 2 we also obtain the analogous result for time-varying population size.

### Proposition 1

(Conditional ancestral process). *The process* 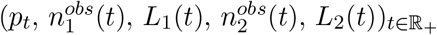 *under ℙ*_**n**_ *is a Markov process with state space* [0, 1] × {0, 1, · · ·, *n*_1_}^2^ × {0, 1, · · ·, *n*_2_}^2^ *described as follows:*

i. 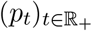 *is a solution to* (1) *with initial density* 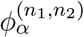. *In particular, it does not depend on the process* 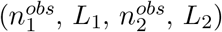.
ii. *The process* 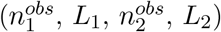 *starts at* (*n*_1_, 0, *n*_2_, 0). *When the current state is* (*p, a*_1_, *ℓ*_1_, *a*_2_, *ℓ*_2_), *this process evolves as*

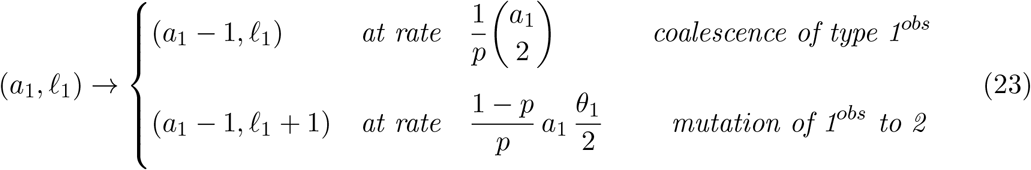

*and, independently*,

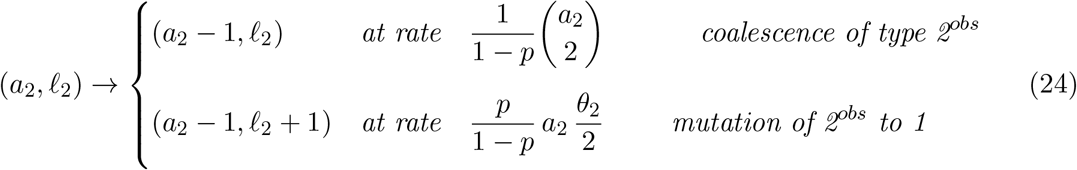

The total rate in (23), at which *a*_1_ decreases by 1, is

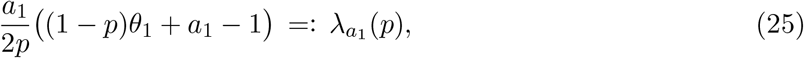

and the one-step transition probabilities are

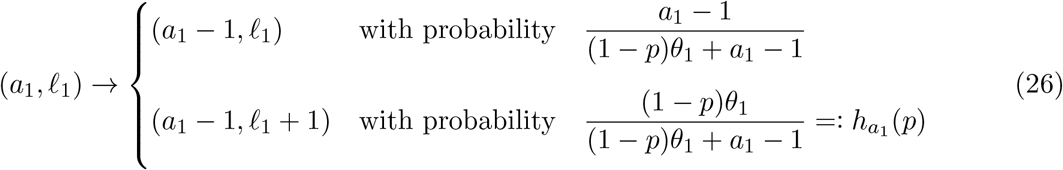

As *t* increases from 0 to ∞, the process 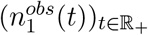 decreases from *n*_1_ to 0, and the process (*L*_1_(*t*))_*t∈ℝ* +_ increases from 0 to a random number *K*_1_ := lim_*t→∞*_ *L*_1_(*t*) ∈ ℕ which is the total number of latent mutations for type 1. Similarly, the total number of latent mutations for type 2 is defined by *K*_2_ := lim_*t→∞*_ *L*_2_(*t*) ∈ ℕ.

We now give a more explicit description of (*K*_1_, *K*_2_) using the frequency process *p* and independent Bernoulli random variables. Note these are conditional on the sample counts (𝒩_1_ = *n*_1_, *𝒩*_2_ = *n*_2_) as in (8).

Let 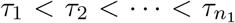 be the jump times of the process 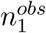. At time *τ*_1_, the process 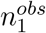 decreases from *n*_1_ to *n*_1_ −1, etc., until finally at *τ*_*n*_, 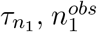 decreases from 1 to 0. It can be checked that 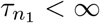 almost surely under ℙ_**n**_ using the ergodicity of the process *p* and (25). Thus 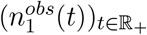 will indeed decrease to 0 eventually under ℙ_**n**_. The allele frequencies at these random times are 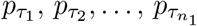. By Proposition 1, under ℙ_**n**_, we have

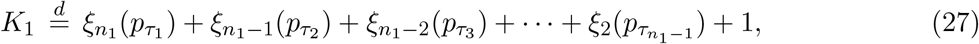

where 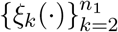 is a family of independent random processes such that, for a constant *p* ∈ [0, 1],

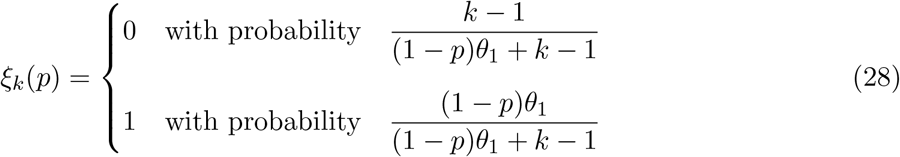

which is a generalization of *ξ*_*k*_ in (7) and (8), where *ξ*_*k*_ ≡ *ξ*_*k*_(0).

Similarly, if we let 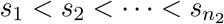 be the jump times of the process 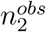, then

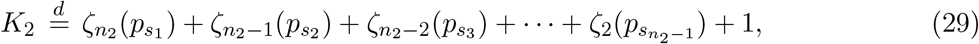

where 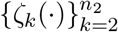 is a family of independent random processes such that, for a constant *p* ∈ [0, 1],

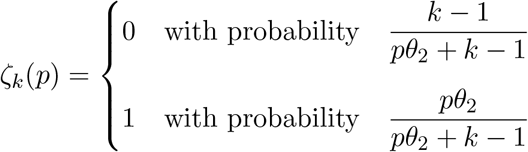

Analogous to *ξ*_*k*_ in reference to *K*_1_, in what follows we will use the notation

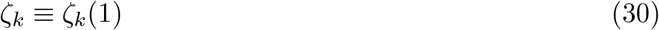

in reference to *K*_2_.

Having described the ancestral process in Proposition 1 and the latent mutations in (27) and (29) under the conditional probability ℙ_**n**_, we study their asymptotic properties under 3 scenarios in the next 3 subsections. These 3 scenarios are a consequence of the asymptotic behaviors of the initial frequency *p*_0_ described in Lemma 1.

### Lemma 4

(Asymptotic initial frequency). *Let n*_1_ ∈ ℕ *be fixed. The initial frequency p*_0_ *converges in probability under* ℙ_**n**_ *to a deterministic constant as follows*.

i. *Suppose n*_2_ ∈ ℕ *is fixed and α* → ∞. *Then p*_0_ → 1.
ii. *Suppose α* ∈ ℝ *is fixed and n*_2_ → ∞. *Then p*_0_ → 0.
iii. *Suppose* 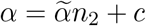 *where*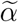, *c* ∈ ℝ *are fixed and n*_2_ → ∞. *Then*

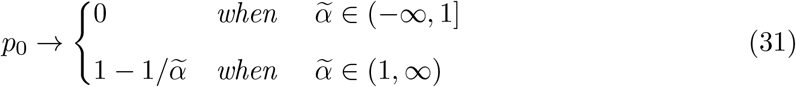

*Furthermore, when* 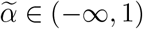, *it holds that n*_2_*p*_0_ *converges in distribution to the Gamma random variable* 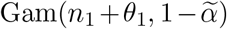 *with probability density function* 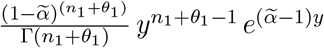.

The proof of Lemma 1 is given in Appendix B.

### 3.1. Scenario (i): strong selection, arbitrary sample size

The first scenario is when |*α*| large with *n*_2_ fixed. We consider the case *α* → +∞ only, since the other case *α* → −∞ follows by switching the roles of type 1 and type 2.

The conditional genealogy of the *n*_1_ + *n*_2_ sampled individuals, under ℙ_**n**_, has three parts with different timescales. First, the *n*_2_ type-2 lineages quickly evolve (coalesce and mutate) as in the Ewens sampling formula, producing *K*_2_ type 1 lineages at a short time *s*_*n*_. Thus, 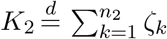 where {*ζ*_*k*_} are independent Bernoulli variables taking values in {0, 1} and having means 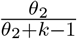. Next, the resulting *n*_1_ + *K*_2_ type 1 lineages will coalesce according to the Kingman coalescent without mutation until only one lineage remains. Hence it takes *O*(1) amount of time for the number of lineages of type 1 to decrease to 1, as *α* → ∞. Finally, it takes a long time, 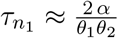, for the single lineage to mutate. In particular, *K*_1_ ≈ 1.

This description is justified by Theorems 1 and 2. See Figure 1 for an illustration.

#### Theorem 1.

*Suppose* (*n*_1_, *n*_2_) ∈ ℕ^2^ *is fixed. Then under* ℙ_**n**_, *as α* → ∞,

i. sup_*t∈*[0,*T*]_ |1 − *p*_*t*_| → 0 *in probability, for any T* ∈ (0, ∞); *and*
ii. *the triplet* 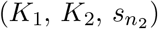 *converges in distribution to* 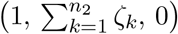, *where* {*ζ*_*k*_} *are independent Bernoulli variables taking values in* {0, 1} *and having means* 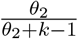.
iii. 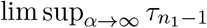 *is stochastically dominated by the height of the Kingman coalescent with n*_1_ + *n*_2_ *leaves*.

*Proof*. By (1), the process *q* := 1 − *p* solves the stochastic differential equation

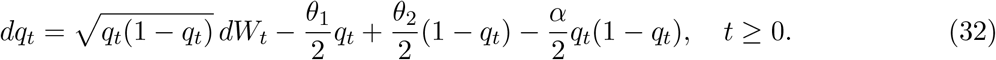

Fix any *ϵ* ∈ (0, 1). We shall to show that ℙ_**n**_(sup_*t∈*[0,*T*]_ *q*_*t*_ *>* 2*ϵ*) → 0 as *α* → ∞.

By the comparison principle (Karatzas and Shreve, 1991, Proposition 2.18 in Chap. 5), we can replace the process *q* by another process 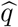 that solves

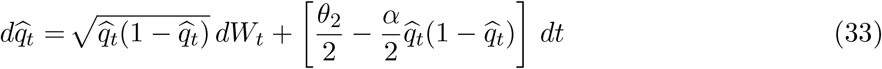

with an initial condition 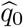 that is equal in distribution to *q*_0_. By Girsanov’s theorem, we can further take away the constant drift 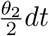. That is, it suffices to show that there exists a probability space (Ω, *ℱ*, ℙ) on which 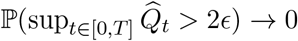 as *α* → ∞, where the process 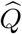 solves

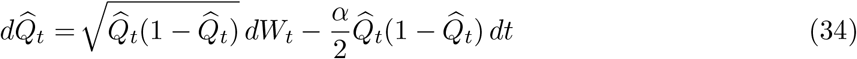

with an initial condition 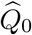 that is equal in distribution to *q*_0_. The initial frequency *q*_0_ → 0 in probability under ℙ_**n**_, by Lemma 1(i). Hence it suffices to show that

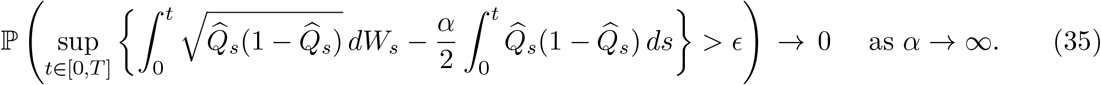

This is true by the time-change representation of the martingale 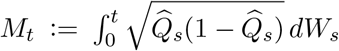 (Karatzas and Shreve, 1991, Theorem 4.6 in Chap. 3) and the fact that

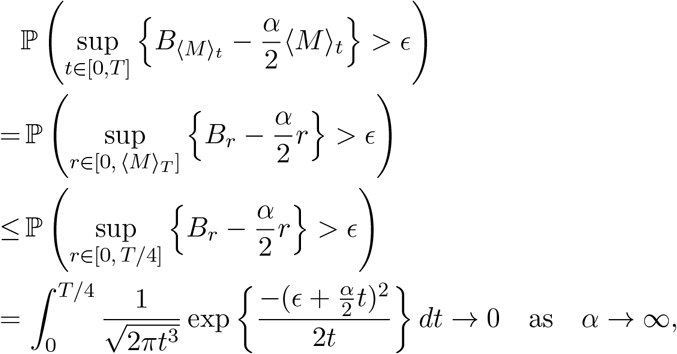

where in the inequality we used the fact that the quadratic variation ⟨*M* ⟩_*T*_ ≤ *T/*4 almost surely (since *q*_*t*_(1 − *q*_*t*_) ≤ 1*/*4 for all *t* ∈ ℝ_+_). Convergence (i) is proved.

Note that the coalescence rate and the mutation rate in (23) converge to *a*_1_(*a*_1_ − 1)*/*2 and 0 respectively as *p* → 1. In (24) both rates converge to infinity but their ratio converges in such a way that the limiting one-step transition probabilities are

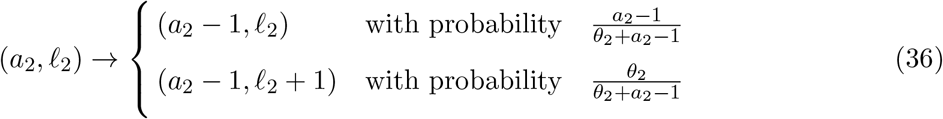

Convergence (ii) then follows from (i) and the representations (27) and (29).

Finally, for part (iii), note that the convergence 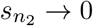 in part (ii) says that the time for type-2 lineages to disappear is negligible. Therefore, it follows from part (i) and (23) that the conditional distribution of 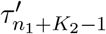, given *K*_2_, converges weakly to the height of the Kingman coalescent with *n*_1_ + *K*_2_ leaves, where 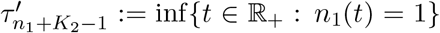 is the first time when the process *n*_1_ decreases to 1. Part (iii) then follows Since 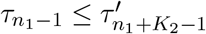 and *K*_2_ ≤ *n*_2_ by definition. □

In Theorem 2 below, we obtain that the mean of the age 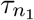 is about 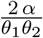.

#### Theorem 2

(Age of the oldest latent mutation of a favorable allele). *Suppose* (*n*_1_, *n*_2_) ∈ ℕ^2^ *is fixed*. *Then under* ℙ_**n**_, *as α* → ∞,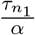 *converges in distribution to an exponential random variable with mean* 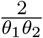. *That is*,

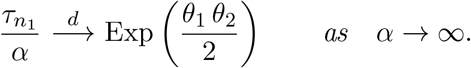

*Proof*. By part (iii) of Theorem 1, it takes *O*(1) amount of time for the number of lineages of type 1 to decrease to 1, as *α* → ∞. It remains to consider the time 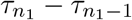 for this single lineage to mutate. Recall the rate of mutation in (23) with *a*_1_ = 1 lineage, for any *t* ∈ ℝ_+_,

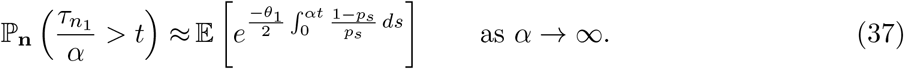

The exponent inside the expectation is, by the ergodic theorem and using the stationary probability density (2) and (3),

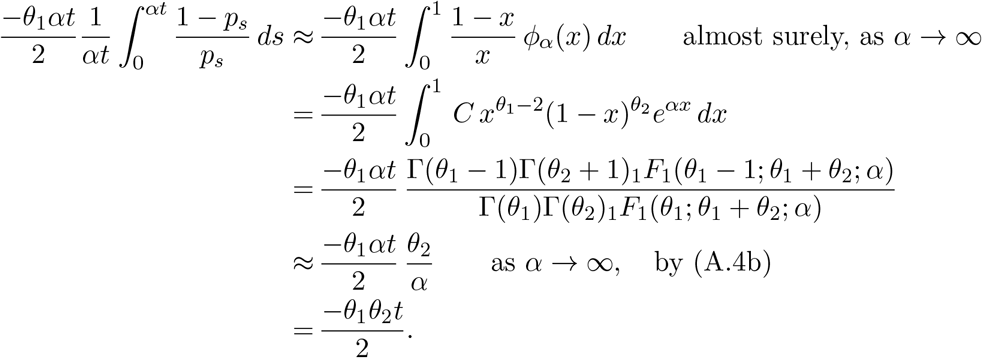

Hence, by (37), 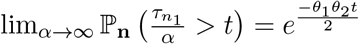 for all *t* ∈ ℝ_+_. The proof is complete. □

In Theorem 2, (37), as in all proofs in this paper, *A* ≈ *B* means that *A/B* → 1 in the limit specified, which is either *α* → ∞ or *n* → ∞. This is equivalent to *A* = *B* + *o*(1) where *B* converges and *o*(1) represents terms which tends to 0 in the limit.

### 3.2 Scenario (ii): arbitrary selection, large sample size

The second scenario is when *n*_2_ large with *α* fixed. We deal with this briefly because it is effectively covered by scenario (iii) when 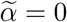.

The conditional genealogy of the *n*_1_ type 1 individuals in the sample can be described as follows. Events among the type 1 lineages occur quickly under ℙ_**n**_ in the sense that 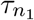 is of order *O*(1*/n*). However, if we measure time in proportion to 1*/n* coalescent time units and measure frequency on the scale of numbers of copies of alleles, then the *n*_1_ type-1 lineages evolve (coalesce and mutate) as in the Ewens sampling formula. In particular, 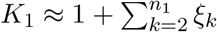, where {*ξ*_*k*_} are independent Bernoulli variables taking values in {0, 1} and having means 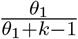.

The rescaled frequency process for type 1 can be described precisely under the rescaling above by the Feller diffusion with drift:

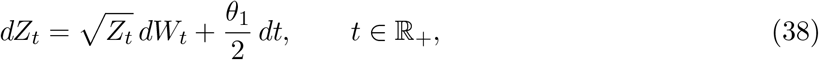

with the initial distribution being the Gamma random variable Gam(*n*_1_ + *θ*_1_, 1). See Figure 2 for an illustration. Remark 1 below explains how this is a special case of scenario (iii), with 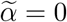.

Equation (38) (also (40) below) is a Cox-Ingersoll-Ross (CIR) model for interest rates in financial mathematics. It has several other names including the Feller process and the square-root process (Dufresne, 2001). It has a unique strong solution. This equation is not explicitly solvable, but its transition density is explicitly known (Vanyolos et al., 2014) and its moments and distributions have been intensively studied.

### 3.3 Scenario (iii): strong selection, large sample size

The third scenario is when both |*α*| and *n*_2_ large with 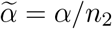 fixed. Lemma 1 implies that

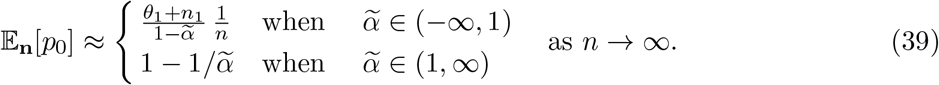

Therefore, it makes sense under this scenario to consider two cases: 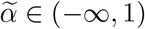 and 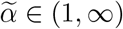.

#### 3.3.1 Case 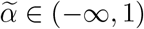

In this case, under ℙ_**n**_ and as *n* → ∞, we have that *p*_0_ = *O*(1*/n*) by Lemma 1. The genealogy of the *n*_1_ type 1 lineages are the same as that in scenario (ii); see Figure 2. This description is justified by Theorems 3-4 below.

Let 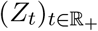 be the ℝ_+_-valued process that has initial state 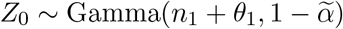 and solves the stochastic differential equation

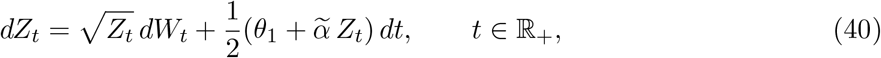

where *W* is the Wiener process.

##### Theorem 3

(Convergence of rescaled genealogy). *Suppose* 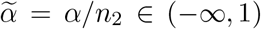 *is fixed. As n* → ∞, *the process* 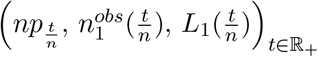 *converges in distribution under* ℙ_**n**_, *in the Skorohod space D*(ℝ_+_, ℝ_+_ × ℤ_+_ × ℤ_+_), *to a Markov process* 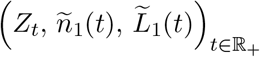 *with state space* ℝ_+_ × {0, 1, ⃛, *n*_1_} × {0, 1, ⃛, *n*_1_} *described as follows:*

i. 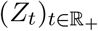 *is a solution to* (40) *with initial state* 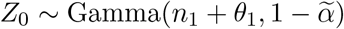. *In particular, its transition kernel does not depend on* 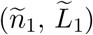.
ii. *Suppose the current state is* (*z, a*_1_, *𝓁*_1_). *Then* 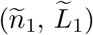 *evolves as*

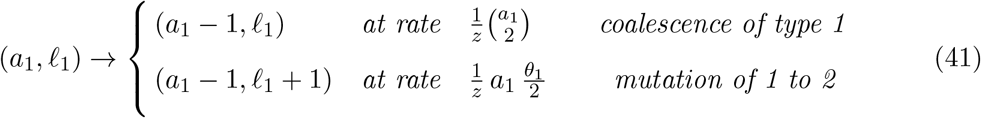

*Proof*. Let *Y*_*t*_ := *np*_*t/n*_. By Lemma 1, under ℙ_**n**_ we have 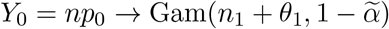. By (1),

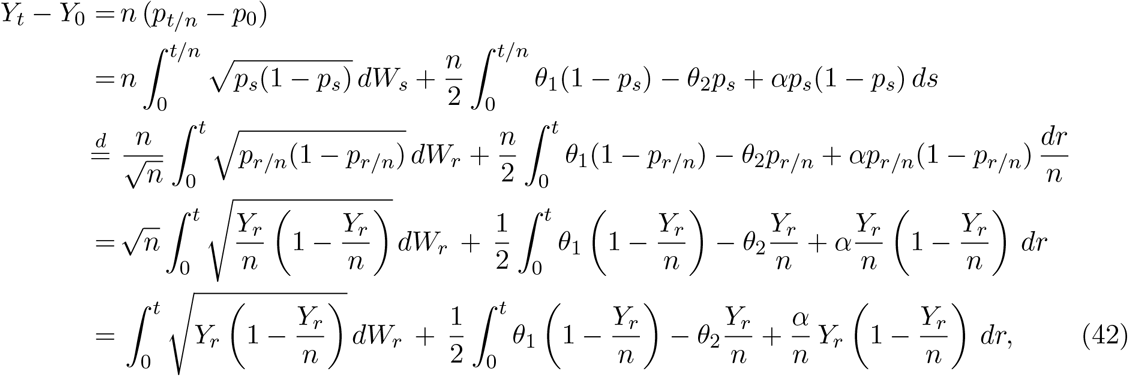

where in the third line above we used the fact that the processes 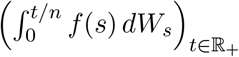 and 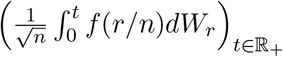 are equal in distribution, where 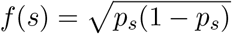.

Using (42), the fact 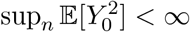 and the assumption 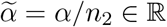 is fixed, we can check by Gronwall’s inequality that lim sup_*n→∞*_ 𝔼_**n**_[sup_*t∈*[0,*T*]_ *Y*_*t*_] *<* ∞ for all *T >* 0. Now, note that equation (40) is the same as (42) after we get rid of the terms 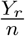 and replace 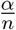 by 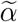. As *n* → ∞, the process (*Y*_*t*_)_*t∈*[0,*T*]_ converges in distribution under ℙ_**n**_ to a process (*Z*_*t*_)_*t∈*[0,*T*]_ with initial state 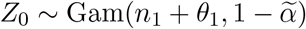 and solving (40).

Using (23), the desired weak convergence in the Skorohod space *D*(ℝ_+_, ℝ_+_ × ℤ_+_ × ℤ_+_) can be checked using a standard compactness argument as in Billingsley (1999, Chap. 2) or Ethier and Kurtz (2005, Chap. 3). That is, we first show that the family is relatively compact: any subsequence has a further subsequence that converges in distribution as *n* → ∞. This can be done using the Prohorov’s theorem. Next, we identify that any subsequential limit is equal in distribution to the process 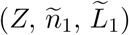, by showing that they solve the same martingale problem. □

By Theorem 3, the jump times of the process 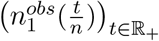 converge to those of the process ñ_1_ as *n* → ∞. See, for instance, Proposition 5.3 in Ethier and Kurtz (2005, Chap. 3). We give a stronger statement and an explicit proof in Theorem 4 below. Theorem 4 also implies that when 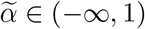, the total number of latent mutations for type 1 is predicted by the Ewens sampling formula, as *n* → ∞. Let 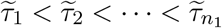 be the jump times of the process *n*_1_ in Theorem 3, at each of which the process decreases by 1.

##### Theorem 4

(Timing of events and number of mutations). *Suppose* 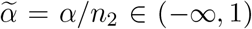 *fixed. Then as n* → ∞,

i. *the random vector* 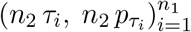 *under* ℙ_**n**_ *converges in distribution to* 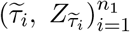.
ii. *K*_1_ *converges in distribution under* ℙ_**n**_ *to* 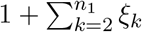,*where* {*ξ*_*k*_} *are independent Bernoulli variables taking values in* {0, 1} *and having means* 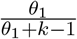.

*Proof*. For part (i), we first give a more explicit description of the jump times 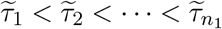, in terms of the function

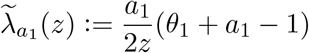

that comes from (41) in Theorem 3. At the first jump time 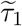, the process *ñ*_1_ decreases from *n*_1_ to *n*_1_ − 1. Thus 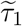 is the first jump time of a Poisson process with time inhomogeneous rate 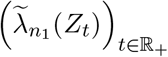, given the trajectory 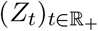. Hence,

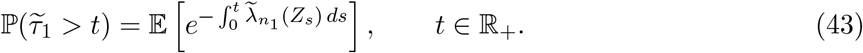

Given 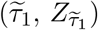, the difference 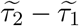 is the first jump time of an independent Poisson process with time inhomogeneous rate 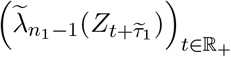. Given 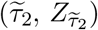, the difference 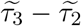 is the first jump time of an independent Poisson process with time inhomogeneous rate 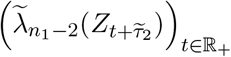 and so on. Finally, given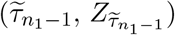, the difference 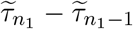 is the first jump time of an independent Poisson process with time inhomogeneous rate 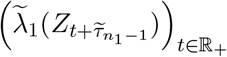.

Using the total rate of type-1 events, 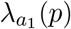 defined in (25), and Theorem 3, as *n* → ∞ we have

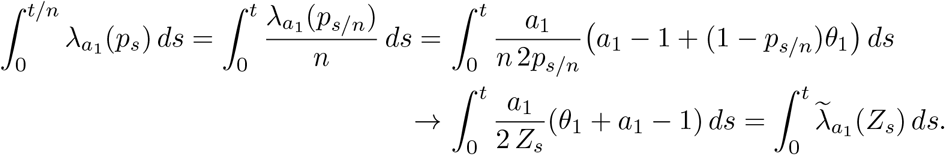

Hence 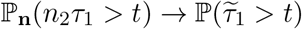 for all *t* ≥ 0, by (43). Combining with Theorem 3, we have that *n*_2_ (*τ*_1_, *p*_*τ* 1_) under ℙ_**n**_ converges in distribution to 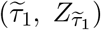 as *n*_2_ → ∞.

Applying the strong Markov property of the proce-ss *Z* at 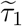 and that of the process *p* at *τ*_*1*_, we can similarly show that *n*_2_ (*τ*_1_, *τ*_2_ − *τ*_1_, *p*_*τ* 1_, *p*_*τ* 2_ − *p*_*τ* 1_) under ℙ_**n**_ convcerges to 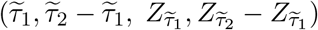 in distribution. Continuing in the same way, we obtain that 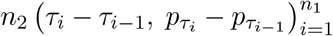 under ℙ_**n**_ converges in distribution to 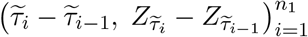, where 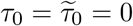. The desired convergence in part (i) then follows.

We now prove part (ii). the vector 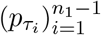 converges in probability to the zero vector in 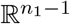 as *n* → ∞, by Theorem 3. This implies that

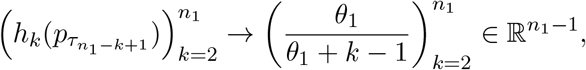

where we recall 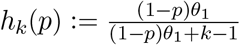 defined in (26). Hence the following weak convergence in 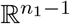 holds:

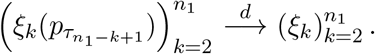

The proof of part (ii) is complete by (27). □

In Wakeley et al. (2023, Appendix), we showed that for the case *α* = 0 (no selection) and *n*_1_ *>* 1, the jump times 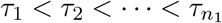 of 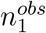 are of order 1*/n* and the re-scaled vector 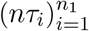 converges in distribution. Theorem 4 therefore generalizes the latter convergence in the presence of selection, in scenario (ii) and in the case 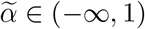 within scenario (iii). This can further be generalized to time-varying populations, as we shall show below. By equation (18) in Wakeley et al.(2023), 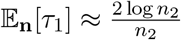 and 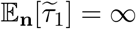 when *n*_1_ = 1 and *α* = 0.

##### Remark 1.

Theorems 3-4 remain valid if “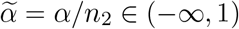 is fixed” is replaced by “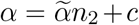 where 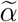, *c* ∈ ℝ are fixed”. In particular, these results hold for scenario (ii).

#### 3.3.2 Case 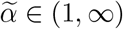

In this case, under ℙ_**n**_ and as *n* → ∞, we have that 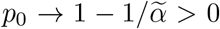 by (31). The process 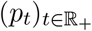 increases very quickly and stays close to 1. As a result, the conditional ancestral process for the *n*_1_ type 1 samples has two parts with different timescales. First it coalesces only as the Kingman coalescent (without mutation) until there is only one lineage. Then it takes a very long time (about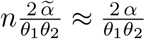) for the single latent mutation to occur. In particular, *K*_1_ ≈ 1.

This description is justified by Theorems 5-6. See Figure 3 for an illustration.

##### Theorem 5.

*Suppose* 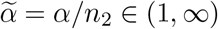 *is fixed. As n* → ∞,*under* ℙ _**n**_,

i. sup_*t∈*[*S,T*]_ |1 − *p*_*t*_| → 0 *in probability, for any* 0 *< S < T <* ∞; *and*
ii. *For any T >* 0, *the process* 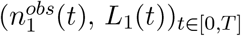 *converges in distribution to a process* 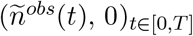, *where* 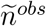 *is a pure death process with jump rate* 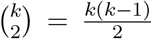.*from k to k* − 1.
iii. *K*_1_ → 1 *in probability*.

*Proof*. We first observe that the process *p* gets close to 1 quickly, when 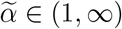, in the sense that for any *ϵ* ∈ (0, 1),

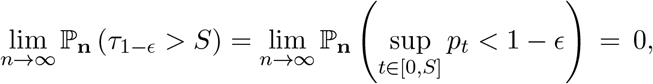

where *τ*_1*−ϵ*_ := inf{*t* ∈ ℝ_+_ : *p*_*t*_ *>* 1 − *ϵ*} is the first time *p* hits a value above 1 − *ϵ*. This is true because 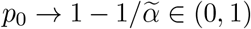 in probability, so that the growth term 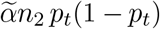 is large at least when *t >* 0 is small. Next, suppose the process *p* starts at 1 − *ϵ* (i.e. the process *q* = 1 − *p* starts at *ϵ*), we show that the exit time of the process *q* out of the interval [0, 2*ϵ*) is longer than *T* with probability tending to 1, as *n* → ∞. More precisely,

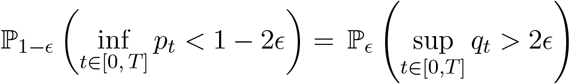

which tends to 0 as *n*_2_ → ∞, as in the proof of Theorem 1(i).

From these two estimates and the strong Markov property of the process *p*, we have that for any *ϵ* ∈ (0, 1),

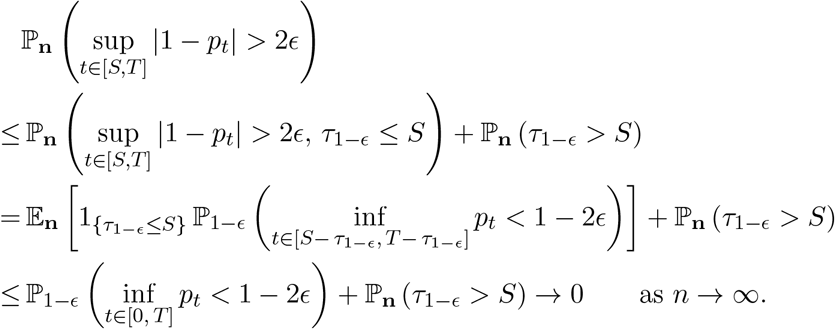

The proof of part (i) is complete.

By part (i) and (25), the times for the type 1 events are of order *O*(1) and *h*_*k*_(*p*_*t*_) → 0 where *h*_*k*_(*p*) is defined in (26). Hence parts (ii) and (iii) follow. □

Now we consider the second part of the genealogy, when there is a single lineage left (i.e. during 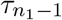 and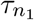).

To estimate the age 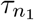 of the single latent mutation, we can ignore the *n*_1_ −1 jump times of the process 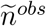 (since they are of order 1 by Theorem 5). The frequency of type 1 is tightly regulated in the sense that it is close to 1 in the sense of Theorem 5(i). However, we need to know “how close it is to 1” in order to get an estimate of 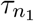, because simply setting *p* = 1 in 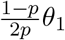 will give us zero.

Theorem 6 below is analogous to Theorem 2. We obtain that the mean of the age 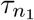 is about 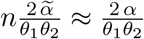 when it is larger than 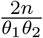 and *n* is large.

##### Theorem 6

(Age of the unique latent mutation). *Suppose* 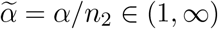 *is fixed. As n* → ∞,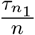 *converges in distribution under* ℙ_**n**_ *to an exponential random variable with mean*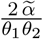. *That is*,

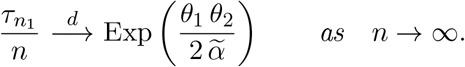

*Proof*. By Theorem 5 (ii), for any *t* ∈ ℝ_+_,

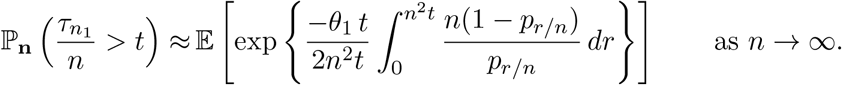

The rest follows exactly as the proof of Theorem 2. □

### 3.4 Time-varying population size

For a population with time-varying size *ρ*(*t*)*N* at forward time *t* where *ρ* is a non-constant function, neither the Moran process nor its diffusion approximation possess a stationary distribution. However, the random background approach of Barton et al. (2004) can be generalized to this setting by considering the time-reversed frequency process.

Our main message in this section is that the limiting results in scenarios (i) and (ii) are robust against continuously-changing population sizes and the initial distribution *µ*_0_ of the initial (ancient) frequency *X*_0_. Roughly speaking, in scenario (ii) as *n*_2_ ∼ *n* → ∞, events among the finite-count *A*_1_ alleles in the sample are so sped up that the population size will have hardly changed by the time all their coalescent and latent mutation events have occurred. The same is true for events among the finite-count *A*_2_ alleles in scenario (i) as *α* → ∞. Events among the finite-count *A*_1_ alleles occur more slowly in scenario (i), but the rate of latent mutations among them remains exceedingly small. The limiting result for scenario (iii) is more subtle. It depends on the large deviation behavior of the present day frequency as *n*_2_ → ∞.

Let *T >* 0 be the present. For comparison with our results for constant population size, we keep the same definitions of *θ*_1_, *θ*_2_ and *α* and we set *ρ*(*T*) = 1. Thus, *N* is the population size at the present time *T*, and *θ*_1_, *θ*_2_ and *α* are the present-day values of these variables. The corresponding values at some other time *t* are *Nρ*(*t*), *θ*_1_*ρ*(*t*), *θ*_2_*ρ*(*t*) and *αρ*(*t*). The demographic function *ρ* could for example represent exponential population growth, in which case *ρ*(*t*) = *ρ*(0)*e*^*βt*^ for some positive constant *β*. This model was used in Wakeley et al. (2023) to illustrate the effects of rapid growth on neutral rare variation in humans. Here we allow that *ρ*(*t*) is piecewise continuous. As will become clear, the key feature of *ρ* for our results is that it is continuous at *T*.

Since the random background approach of Barton et al. (2004) was formulated based on the lineage dynamics of the Moran model, we begin by describing the diffusion process arising from a Moran model with time-varying population size.

#### Lemma 5

(Diffusion limit for time-varying Moran model). *Let ρ* : ℝ_+_ → (0, ∞) *be a piecewise continous function with finitely many jumps, and N be a positive integer. Consider the discrete-time Moran process in which, at step k* = [*N* (*N* − 1)*t/*2], *the total population size is* [*ρ*(*t*)*N*] *and N is replaced by ρ*(*t*)*N in the one-step transition probabilities* (13)*-*(14). *Suppose u*_1_ = *θ*_1_*/N, u*_2_ = *θ*_2_*/N and s* = −*α/N. Then as N* → ∞, *the relative frequency of A*_1_ *at step* [*N* (*N* − 1)*t/*2] *converges in distribution to X*_*t*_ *solving*

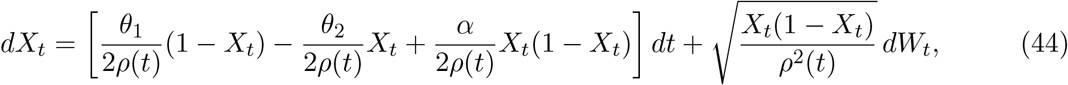

*where W*_*t*_ *is the Wiener process, provided that the initial relative frequency converges to X*(0).

Setting *ρ*(*t*) ≡ 1 for all *t* ∈ ℝ_+_, or *β* = 0 in the exponential growth model, makes (44) identical to (1). The term *ρ*^*2*^ (*t*) in the denominator inside the square root comes from the diffusion timescale of the Moran model: for a population of constant size, one unit of time in the diffusion is *N* (*N* − 1)*/*2 ∝ *N* ^2^ time steps in the discrete model. To explain the term 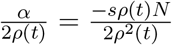, note that the rate of change of *X*_*t*_ due to selection is proportional to the product of the total size *ρ*(*t*)*N* and the parameter *s*, which is then multiplied by 1*/ρ*^2^(*t*) because the timescale in (44) is defined in terms of the present-day population size *N*. The proof of Lemma 1 is given in Appendix B.

#### Remark 2

(Wright-Fisher model with varying size). The analogous diffusion process *X*^WF^ for the discrete Wright-Fisher model with total size [*ρ*(*t*)*N*] in generation [*Nt*] is *different from* the process *X* in (44), except in the case *ρ*(*t*) ≡ 1 for all *t* ∈ ℝ_+_. This diffusion solves the SDE

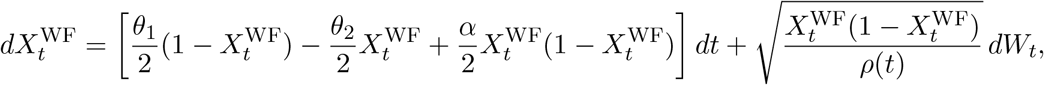

which is the adaptation of equation (1) in Schraiber et al. (2016) to our haploid model of selection and recurrent mutation; see also equation (21) in Evans et al. (2007). The generators of *X*^WF^ and *X* are related by 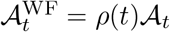 for all *t* ∈ ℝ_+_. In other words, the diffusion *X*^WF^ from the discrete Wright-Fisher model is sped up by the factor *ρ*(*t*) at time *t*.

To compare *X* and *X*^WF^, we can perform deterministic time-changes to normalize their diffusion coefficients to be the same. In general, suppose *X* satisfies the SDE *dX*_*t*_ = *b*(*t, X*_*t*_)*dt* + *σ*(*t, X*_*t*_) *dW*_*t*_ and *Y* is a time-change of *X* defined by *Y*_*r*_ := *X*_*ψ*(*r*)_, where *ψ* is any fixed continuous and strictly increasing function, then

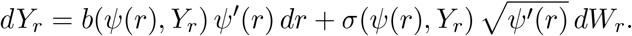

Hence, when *X* is a (weak) solution to (44) and *ψ* = *g*^*−*1^ where *g* is the unique continuous function such that 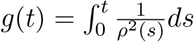, we obtain that 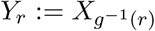 solves

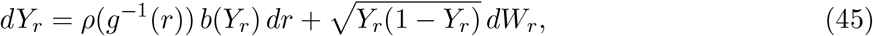

where 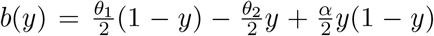. Analogously, following Schraiber et al. (2016)—see their equation (6) and the SDE below it—and taking *f* such that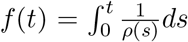, we find that 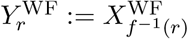 solves

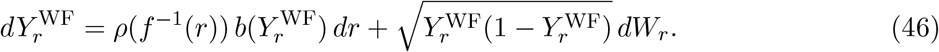

Since *f* ≠ *g* unless *ρ*(*t*) ≡ 1 for all *t* ∈ ℝ_+_, we have that *Y* ≠*Y* ^WF^, i.e. 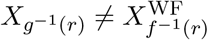 in general. Nonetheless, (45) and (46) have the same form, the only difference being the way time *r* in these diffusions is related back to time *t* in the discrete models.

Note that the law of the present-time frequency of *A*_1_ in the model of Lemma 1 depends on the distribution *µ*_0_ of the initial frequency *X*(0). This law is denoted by ℙ_*µ* 0_ (*X*_*T*_ ∈ *dy*).

Suppose a sample of *n* individuals are picked uniformly at random at the present time *T >* 0, i.e. when the frequency of *A*_1_ is *X*_*T*_, and we know that *n*_1_ of them are of type 1 (and *n* − *n*_1_ are of type 2). Let 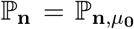 be the conditional probability measure given the sample count **n** = (*n*_1_, *n*_2_). We also denote the conditional law of the present frequency *p*_0_ = *X*_*T*_, under ℙ_**n**_, by ℒ^**n**^ := ℙ_**n**_(*X*_*T*_ ∈ *dy*) = ℙ_*µ*0_ (X_T_ ∈ *dy* | **n**). Then

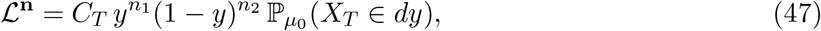

where 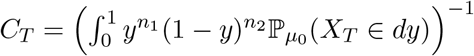 is a normalizing constant. This follows from Bayes’ theorem, just like (5) did, but with prior distribution ℙ_*µ* 0_ (*X*_*T*_ ∈ *dy*).

Similar to Proposition 1, the conditional ancestral process in the diffusion limit can be described as follows. This description involves the backward frequency process

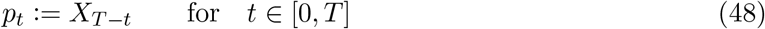

which is by definition the time-reversal of the process *X*.

#### Proposition 2

(Conditional ancestral process). *Let T >* 0 *and µ*_0_ ∈ 𝒫 ([0, 1]) *be fixed, and the demographic function ρ be as in Lemma 1*. *The process* (*p*_*t*_, 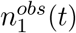, *L*_1_(*t*), 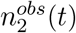, *L*_2_(*t*))_*t∈*[0,*T*]_ *under* ℙ_**n**_ *is a time-inhomogeneous Markov process with state space* [0, 1] × {0, 1, ⃛, *n*_1_}^2^ × {0, 1, ⃛, *n*_2_}^2^ *described as follows:*

i. (*p*_*t*_)_*t∈*[0,*T*]_, *defined by* (48), *has the law of* (*X*_*T −t*_)_*t∈*[0,*T*]_ *under* ℙ_**n**_. *In particular, it has initial distribution ℒ*_**n**_ *in* (47) *and it does not depend on the process* 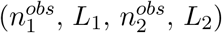.
ii. *The process* 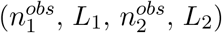 *starts at* (*n*_1_, 0, *n*_2_, 0). *When this process is at time t and the current state is* (*p, a*_1_, *𝓁*_1_, *a*_2_, *𝓁*_2_), *this process evolves as*

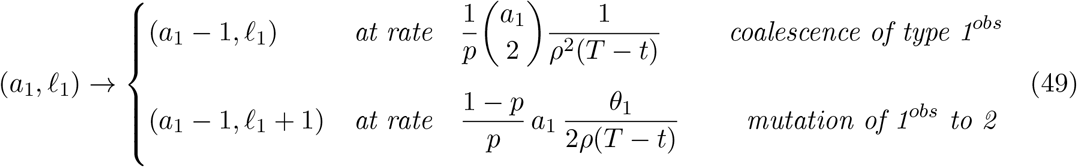

*and, independently*,

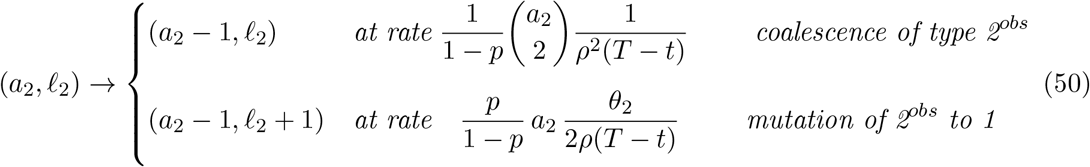

Note the term *ρ*(*T* − *t*) in (49)-(50) indicates the dependence of the conditional ancestral process on the demographic function. Nonetheless, Proposition 2 still gives a description for *K*_1_ and *K*_2_ in terms of Bernoulli random variables, like (27) and (29) respectively. For example, under ℙ_**n**_, (27) still holds but (28) needs to be modified. Indeed, given 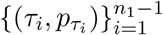, the random variables 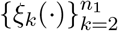 are independent and

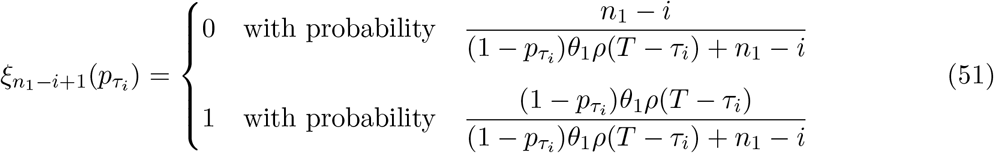

which further generalizes *ξ*_*k*_(*p*) in (28) to include *ρ*(*t*). An analogous description holds for *K*_2_.

#### Remark 3.

A more explicit description for the process (*X*_*t*_)_*t∈*[0,*T*]_ under ℙ_**n**_, hence also that for its time-reversal (*p*_*t*_)_*t∈*[0,*T*]_, can be obtained by Doob’s h-transform (Doob, 1957, 2001). More precisely, we define the function

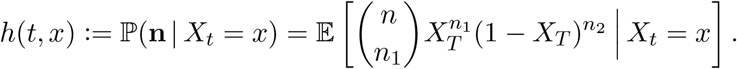

Then (*X*_*t*_)_*t∈*[0,*T*]_ under the conditional probability ℙ_**n**_ solves the SDE

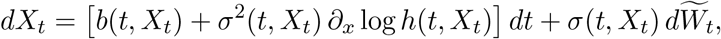

where 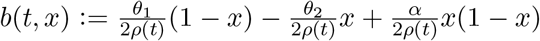 and 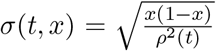 are the coefficients in (44), and 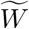 is a Brownian motion. Sufficient conditions on the function *ρ* for which the process (*p*_*t*_)_*t∈*[0,*T*]_ satisfies a stochastic differential equation may be deduced from an integration by parts argument as in Millet et al. (1989).

Next, we look at asymptotics. The following analogue of Lemma 1 holds for any initial distribution *µ*_0_ of *X*(0) and any demographic function *ρ* that is bounded and positive. Note that in Lemma 1, *µ*_0_ = *ϕ*_*α*_ depends on *α*, but here *µ*_0_ is fixed.

#### Proposition 3.

*Let T >* 0 *and µ*_0_ ∈ 𝒫 ([0, 1]) *be fixed, and the demographic function ρ be as in Lemma 1*. *The following convergences in 𝒫*([0, 1]) *hold*.

i. *Suppose n*_2_ *is fixed and α* → ∞. *Then ℒ*_**n**_ → *δ*_1_.
ii. *Suppose α is fixed and n*_2_ → ∞. *Then ℒ*_**n**_ → *δ*_0_.
iii. *Suppose*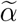, *c* ∈ ℝ *are fixed and* 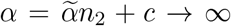. *Suppose* ℙ_*µ* 0_ (*X*_*T*_ ∈ *dy*) *has a density p*(*T, µ*_0_, *y*) *dy and there exists a large deviation rate function ℐ* : [0, 1] → [0, ∞] *such that for each y* ∈ [0, 1],

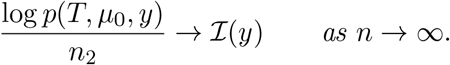

*Suppose also that* (1 − *y*) *e* ^*ℐ*(*y*)^ *has a unique maximum at y*_*\**_ ∈ [0, 1]. *Then* 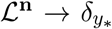, *and* 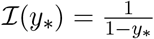.

#### Remark 4.

The assumptions in (iii) hold when *ρ*(*t*) ≡ 1 and *µ*_0_ = *ϕ*_*α*_ in (2), i.e. constant population size with stationary initial condition. In this case, 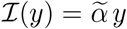 is the linear function, and 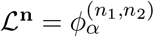 in (6). The rate function has a phase transition at 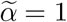, as shown in Lemma 1. Namely, *y*_*\**_ = 0 when 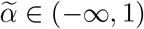 and 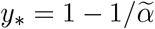 when 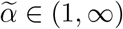.

#### Remark 5.

The large deviation principle for ℙ_*µ* 0_ (*X*_*T*_ ∈ *dy*) as *n*_2_ → ∞ can be checked using the Gärtner-Ellis theorem (Dembo and Zeitouni, 2009, Theorem 2.3.6). When it holds, the rate function ℐis equal to the Legendre transform of the function

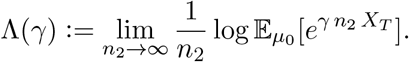

*Proof*. A proof follows from that of Lemma 1. Let *f* ∈ *C*_*b*_([0, 1]), a bounded continuous function on [0, 1]. Then

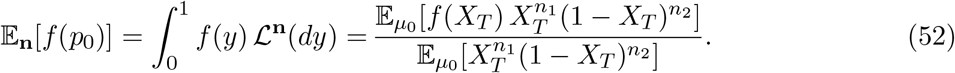

For part (i), note that if (*n*_1_, *n*_2_) is fixed and *α* → ∞, then ℙ_*µ* 0_ (*X*_*T*_ *<* 1 − *ϵ*) → 0 for any *ϵ >* 0 as in the proof of Theorem 5(i). Hence 𝔼_*µ* 0_ [|*f* (*X*_*T*_) − *f* (1)|] → 0 for any *f* ∈ *C*_*b*_([0, 1]). In particular, ℙ_*µ*_ (*X*_*T*_ ∈ *dy*) → *δ*_1_. Hence ℒ_**n**_ tends to *δ*_1_ in 𝒫 ([0, 1]), as *α* → ∞.

For part (ii), note that (1 − *y*)^*n*2^ has maximum at *y* = 0, and *y*^*n*1^ ℙ_*µ*_ (*X*_*T*_ ∈ *dy*) does not depend on *n*_2_. Hence ℒ_**n**_ tends to *δ*_0_ in 𝒫 ([0, 1]), as *n*_2_ → ∞.

For part (iii), the numerator of (52) is

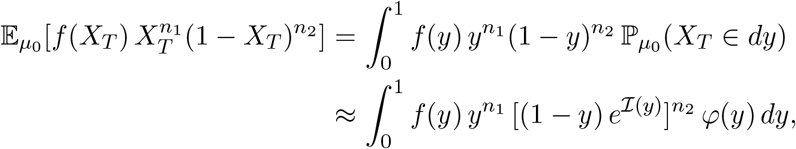

for some function *φ* such that 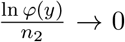, by assumptions of part (iii). Since (1 − *y*) *e* ^*ℐ*(*y*)^ has a unique maximum at *y*_*\**_, lim_*n→∞*_ 𝔼_**n**_[*f* (*p*_0_)] = *f* (*y*_*\**_) by (52) and a standard argument as in the proof of Lemma 1. □

By Proposition 2 and Proposition 3, similar limiting results for the conditional coalescent process for scenarios (i) and (ii) hold for any positive function *ρ* : [0, *T*] → (0, ∞) that is *continuous near the current time T*. Note that *ρ* is bounded away from zero on any compact time interval [0, *T*] and therefore analogous approximations for the frequency process (*p*_*t*_)_*t∈*[0,*T*]_ under ℙ_**n**_ still hold, where the new approximating functions now involve the *h* function in Remark 3.

More precisely, in scenario (i), Theorem 1 still holds, and Theorem 2 still holds but with a possibly different limiting random variable. Hence 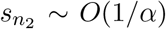 is very small and 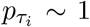, so *K*_1_ → 1 by (51), and the single latent mutation for type 1 is very old.

In scenario (ii), 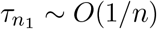 is very small, and recalling that *ρ*(*T*) = 1, we have 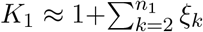 by (51), where {*ξ*_*k*_} are independent Bernoulli variables taking values in {0, 1} and having means 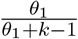. Theorem 3 with *α* ∈ ℝ fixed still holds, but the statement needs to be modified because the approximating process *Z* in (40) will be replaced by another one that involves the *h* function in Remark 3.

Scenario (iii) is harder to analyze and we leave it for future work. We conjecture that if *y*_*\**_ = 0, then the conditional genealogy behaves like scenario (ii); and if *y*_*\**_ ∈ (0, 1], then the conditional genealogy behaves like scenario (i).

## 4. Conditional ancestral selection graph

Our aim in this section is to see how the results of the previous section can be obtained from a different model: the ancestral selection graph. We concentrate on the ancestry of focal allele *A*_1_ and on constant population size, and proceed more heuristically than in the previous section.

The ancestral selection graph is an augmented coalescent model for the joint distribution of the gene genealogy and the allelic states of the sample (Krone and Neuhauser, 1997; Neuhauser and Krone, 1997). It includes the usual coalescent rate 1 per pair of lineages and mutation rate *θ/*2 per lineage. Additionally, under the stationary model of Section 1, it includes a *branching* rate of |*α*|*/*2 per lineage. When a branching event occurs, the lineage splits into an *incoming* lineage and a *continuing* lineage. One of these is *real*, meaning it is included in the gene genealogy. The other is *virtual*, meaning it is there only to model the gene genealogy correctly with selection. Which is which could be resolved if their allelic states were known: the incoming lineage is real if its allelic type is the one favored by selection, otherwise the continuing lineage is real. But the allelic states are not known in the construction of the ancestral selection graph.

The conditional ancestral selection graph models gene genealogies given a sample with allelic states specified (Slade, 2000a,b; Fearnhead, 2001, 2002; Stephens and Donnelly, 2003; Baake and Bialowons, 2008). In this case it is known which lineages are real and which are virtual. This allows a simplification in which there is a reduced rate of branching and only virtual lineages of the disfavored type are produced (Slade, 2000a). A second simplification is possible if mutation is parent-independent: then any lineage which mutates may be discarded (Fearnhead, 2002).

We assume parent-independent mutation, specifically *θ*_1_ = *θπ*_1_ and *θ*_2_ = *θπ*_2_, with *π*_1_ + *π*_2_ = 1. Any two-allele mutation model can be restated in this way, leaving the stationary probability density (2) and the sampling probability (4) unchanged. But doing so introduces “spurious mutations to one’s own type” (Donnelly, 1986) or “empty mutations” (Baake and Bialowons, 2008) which occur only in the model and do not correspond to a biological process. These are not latent mutations. Including them allows us to discard real *A*_2_ lineages and any virtual lineage once these mutate, but we must distinguish between empty and actual mutations in the ancestry of *A*_1_.

The resulting conditional process tracks the numbers of real and virtual ancestral lineages from the present time *t* = 0 back into the past. Let *r*_1_(*t*), *r*_2_(*t*) and *v*_*i*_(*t*), where *i* = 1 if *α <* 0 or *i* = 2 if *α >* 0, be the numbers of real type-1, real type-2 and virtual type-*i* lineages at past time *t*. The process begins in state *r*_1_(0) = *n*_1_, *r*_2_(0) = *n*_2_, *v*_*i*_(0) = 0 and stops when *r*_1_(*t*) + *r*_2_(*t*) = 1. We suppress *t* in what follows, and focus on the instantaneous transition rates of the process.

The conditional ancestral process is obtained by considering rates of events in the unconditional process, which has total rate (*r*_1_ + *r*_2_ + *v*_*i*_)(*θ* +|*α*| + *r*_1_ + *r*_2_ + *v*_*i*_ − 1)*/*2, then weighting rates of events depending on how likely they are to produce the sample. Rates of some events are down-weighted to zero. For instance, the sample could not have been obtained if there were a coalescent event between lineages with different allelic types, whereas in the unconditional process these happen with rate *r*_1_*r*_2_ plus either *r*_2_*v*_1_ or *r*_1_*v*_2_, depending on whether *α <* 0 or *α >* 0.

Rates of events for which the sample has a non-zero chance of being observed are up-weighted or down-weighted by ratios of sampling probabilities like (4). This method of conditioning a Markov process on its eventual outcome is stated simply in Kemeny and Snell (1960, p. 64), a familiar example being the Wright-Fisher diffusion conditioned on eventual fixation (Ewens, 2004, p. 89), and is characterized more generally by Doob’s h-transform (Doob, 1957, 2001). In the conditional ancestral selection graph, the Markov process is the (unconditional) ancestral process of Krone and Neuhauser (1997) and the eventual outcome is the sample with allelic states specified.

In our formulation, the samples and their ancestral lineages all are distinguishable, which we denote with a subscript “o” for ordered as in Wakeley et al. (2023). The probability of any particular allelic configuration in the ancestry of the sample, in which there are *r*_1_ lineages of type 1, *r*_2_ lineages of type 2 and *v*_*i*_ lineages of type *i* ∈ {1, 2}, is

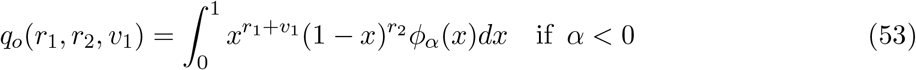

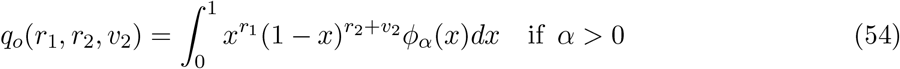

with *ϕ*_*α*_(*x*) as in (2). Note, the additional binomial coefficient in the sampling probability (4) is the number of possible orderings of a sample containing *n*_1_ and *n*_2_ copies of *A*_1_ and *A*_2_.

The rate of any particular event with non-zero probability in the conditional process is the product of its rate in the unconditional process and a ratio of sampling probabilities from either (53) or (54). For event 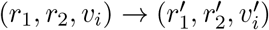, the required ratio is 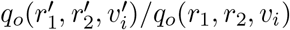. The denominator *q*_*o*_(*r*_1_, *r*_2_, *v*_*i*_) is the probability of the sample given all events so far in the conditional ancestral process, which have led to the current state (*r*_1_, *r*_2_, *v*_*i*_), and the numerator 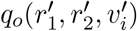 is the probability of the sample given these events and the event 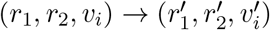. Appendix C provides the details of how the minimal ancestral process we use here to model latent mutations in the ancestry of the sampled copies of allele *A*_1_ is obtained from the full conditional ancestral process, using the simplifications of Slade (2000a) and Fearnhead (2002).

The resulting conditional ancestral process differs depending on whether *α <* 0 or *α >* 0 but in either case it includes five possible transitions from state (*r*_1_, *r*_2_, *v*_*i*_). If *α <* 0,

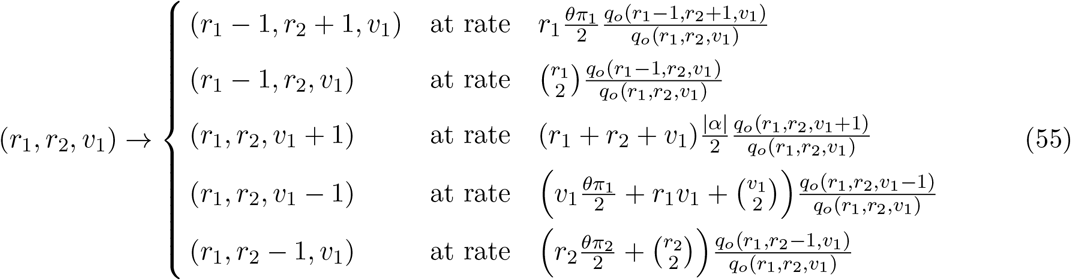

whereas if *α >* 0,

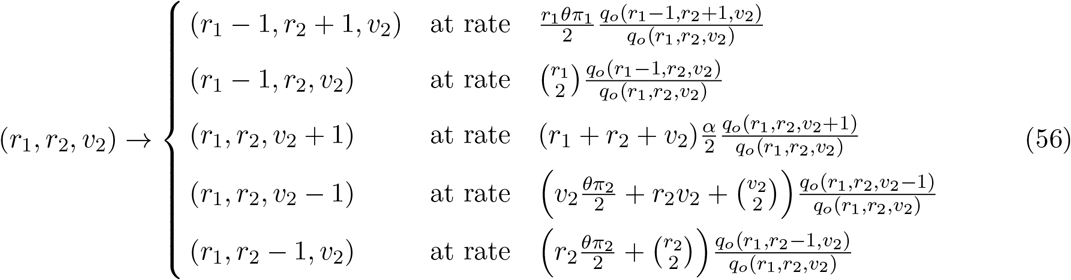

which differ owing to the different resolutions of branching events when *α <* 0 versus *α >* 0. We may note that the total rates of events in (55) and (56) are less than in the unconditional ancestral process because the conditional process has a reduced rate of branching (Slade, 2000a) and because empty mutations do not change the number or types of ancestral lineages. If *α <* 0, the total rate is *r*_1_*θπ*_1_*/*2 + *r*_2_|*α*|*/*2 less, whereas if *α >* 0, it is *r*_1_*θπ*_1_*/*2 + *r*_1_*α/*2 less.

Asymptotic approximations for the ratios 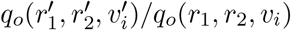 in these rates of events can be obtained using the results in Appendix A. In the following three subsections we present approximations to the conditional ancestral process for our three scenarios of interest: (i) |*α*| large with *n*_2_ fixed, (ii) *n*_2_ large with *α* fixed, and (iii) both |*α*| and *n*_2_ large with 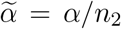 fixed.

Because initially *r*_2_ = *n*_2_, we consider *r*_2_ large in the scenarios with *n*_2_ large. For each scenario, we compute the transition rates up to leading order in |*α*| or *r*_2_, then consider how these conform to the corresponding results of Section 3.

### 4.1. Scenario (i): strong selection, arbitrary sample size

Here |*α*| is large with *n*_2_ fixed, along with *n*_1_ and *θ*. In Section 3.1, Theorem 1, we treated the ancestries of *A*_1_ and *A*_2_ simultaneously as *α* → +∞, so that *A*_1_ was favored and *A*_2_ was disfavored. Here we cover these same two possibilities by modeling the ancestry of *A*_1_, using (55) when *A*_1_ is disfavored (*α <* 0) and (56) when *A*_1_ is favored (*α >* 0). We disregard the ancestry of the non-focal allele *A*_2_ except insofar as it is needed to model events in the ancestry of *A*_1_.

When *α <* 0, using (A.4a) in (55) gives

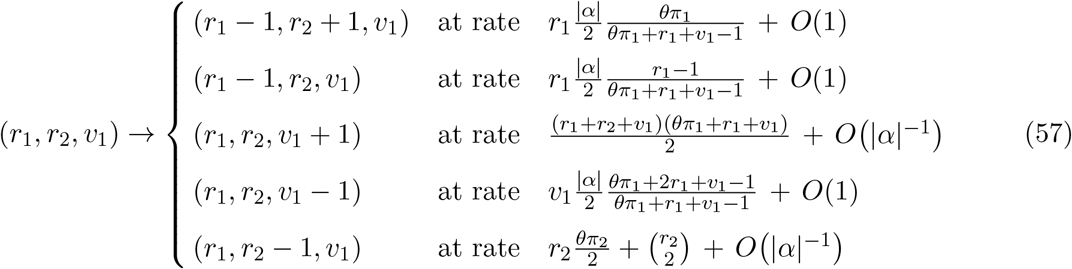

for the ancestry of a disfavored allele under strong selection (as *α* → −∞). Latent mutations and coalescent events occur with rates proportional to |*α*|. Virtual lineages are removed similarly quickly but are produced at a much lower rate. So *v*_1_ will stay zero during the *O*(1*/*|*α*|) time it takes for the requisite *n*_1_ latent mutations or coalescent events to occur. Then the analogous result to (36), namely (7) and (8), follows from the first two lines of (57). Coalescence and mutation among the copies of *A*_2_ occur at the slower rate, so none of these should occur before all the type-1 lineages disappear. These results were first suggested in Wakeley (2008).

When *α >* 0, using (A.4b) in (56) gives

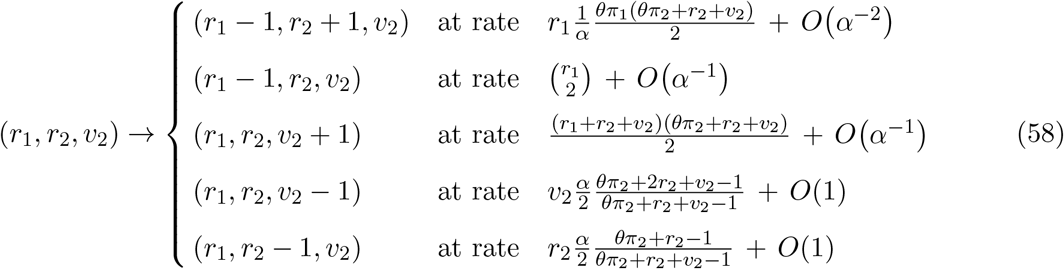

for the ancestry of a favored allele under strong selection (as *α* → +∞). Now *A*_2_ is undergoing the fast process just described for *A*_1_ in (57), so these lineages will disappear quickly. Again the rate of removal of virtual lineages greatly exceeds their rate of production. In *O*(1*/α*) time, the ancestral state will become (*r*_1_, *r*_2_, *v*_2_) = (*n*_1_, 0, 0). But now with *A*_1_ favored, the rates of coalescence and latent mutation differ by a factor of *α*, so the first *n*_1_ − 1 events will be coalescent events, followed by a long wait for a single latent mutation with rate *θ*^2^*π*_1_*π*_2_*/*(2*α*) as in Theorem 2.

### 4.2 Scenario (ii): arbitrary selection, large sample size

Here *n*_2_ is large with *α* fixed, along with *n*_1_ and *θ*. Because *r*_2_ = *n*_2_ at the start of the ancestral process, we present rates of events to leading order in 1*/r*_2_. In Section 3.2 we deferred this scenario to Section 3.3, because in the limit it is equivalent to 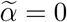. Of course, there are two ways for 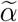 to approach zero, and the sign of 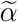 matters in (40) for any 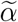 not strictly equal to zero. Here we consider the two cases, *α <* 0 and *α >* 0, separately.

When *α <* 0, using (A.5) in (55) gives

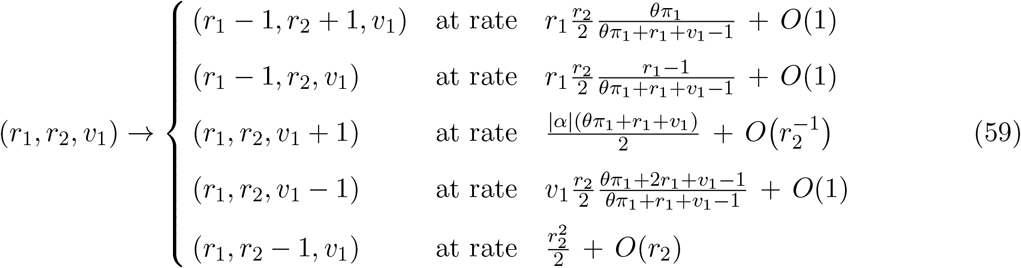

This differs from the neutral case (Wakeley et al., 2023) only by the possibility of virtual lineages. As in (57), these will be removed quickly if they are produced. The process of latent mutation and coalescence happens in *O*(1*/r*_2_) time, with relative rates in the first two lines of (59) again giving (7) and (8). Because *r*_2_ → ∞, this approximation will hold long enough for the required fixed number of events among the *A*_1_ lineages to occur, despite the rapid decrease of *r*_2_ in the last line of (59). A proof of this is given in Wakeley et al. (2023, Appendix). Theorem 4 addresses the corresponding issues for the model of Section 3.

When *α >* 0, using (A.5) in (56) gives

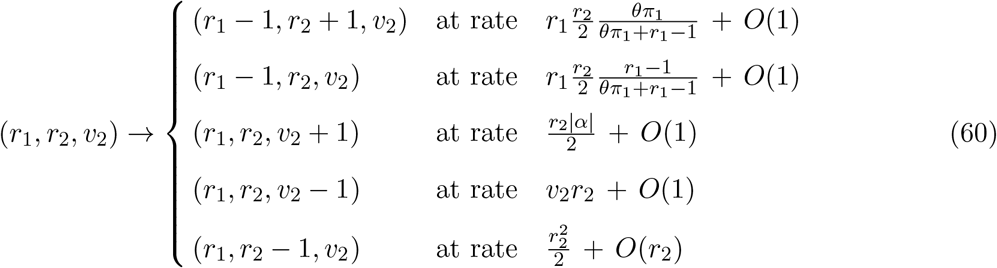

which differs from (59) in two ways. Now the rate of production of virtual lines is non-negligible. But here their presence does not affect the rates of latent mutation and coalescence. Again we have (7) and (8), and the process of latent mutation and coalescence happens in *O*(1*/r*_2_) time.

### 4.3 Scenario (iii): strong selection, large sample size

Here both |*α*| and *n*_2_ are large with 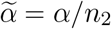 fixed, along with *n*_1_ and *θ*. Again since the process begins with *r*_2_ = *n*_2_, we present rates of events to leading order in 1*/r*_2_. Because the conditional ancestral process differs for *α <* 0 versus *α >* 0, i.e. with (55) and (56), and the asymptotic approximation we use for the hypergeometric function differs for 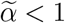 versus 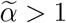, i.e. with (A.6a) and (A.6b), here we have three cases. Note these are the same three cases in (11a), (11b) and (11c).

When 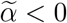, using (A.6a) in (55) gives

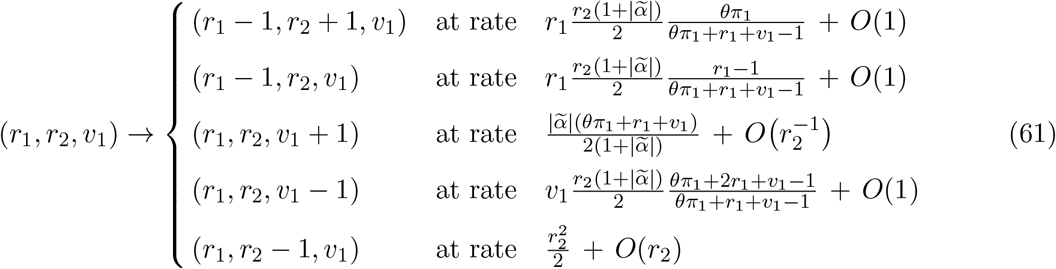

which is comparable to (57) and (59). Again we may effectively ignore virtual lineages. The rates of latent mutation and coalescence in (57) and (59) differ only by the interchange of *r*_2_ for |*α*|. In (61), the factor 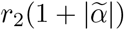 encompasses the effects of both. The larger 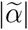 is, the more quickly these events will occur, and again (7) and (8) describe the number of latent mutations.

When 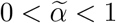, using (A.6a) in (56) gives

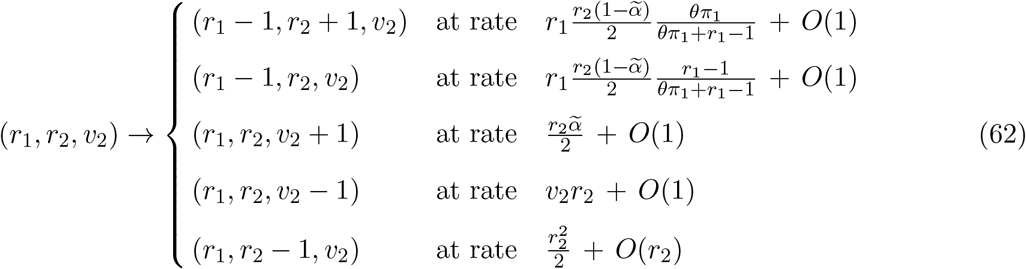

which is comparable to (60). In contrast to (61), now with *A*_1_ favored, the larger 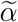 is (i.e. the closer it is to 1) the smaller the rates of latent mutation and coalescence become. Otherwise, for any given 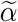, the same conclusions regarding latent mutations and their timing follow from (62) as from (61), and these conform to what is stated in Theorem 4.

When 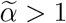, using (A.6b) in (56) gives

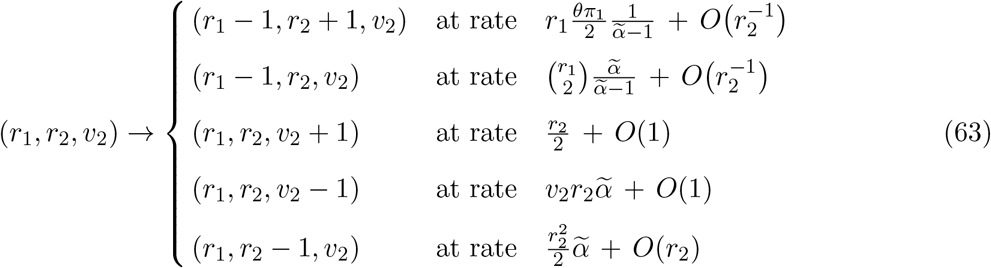

which paints a very different picture. Whereas (57), (59), (60), (61) and (62) all give the Ewens sampling result described by (7) and (8) and have these events occurring quickly on the coalescent time scale, (63) is rather like (58) in that the rates of latent mutation and coalescence are too slow to register on the time scale of events involving the non-focal allele *A*_2_. The overwhelmingly most frequent events in (63) will be coalescent events between *A*_2_ lineages at rate 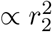, so an effectively instantaneous transition will occur from *r*_2_ large to *r*_2_ comparable to *r*_1_. Then this case (63) will collapse quickly to the corresponding case (58) where coalescence without mutation will happen among the *A*_1_ followed by a long wait for a single latent mutation. For the model in Section 3.3.2, this is described by Theorem 5 and Theorem 6. Finally we may note that initially the rates of latent mutation and coalescence in (63) are precisely those predicted for the model in Section 3.3.2 from (23) starting at 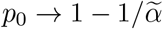 as specified for 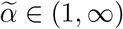 in (31).

## 5. Discussion

In this paper, we have considered a two allele model at a single genetic locus subject to recurrent mutation and selection in a large haploid population with possibly time-varying size. We assumed that a sample of size *n* was drawn uniformly from an infinite population under the diffusion approximation. By extending the framework of Barton et al. (2004), we described the asymptotic behaviors of the conditional genealogy and the number of latent mutations of the sample, given the sample frequencies of the two alleles. This moves beyond what is in Wakeley et al. (2023) by the inclusion of selection and by the use of an entirely different model, i.e. coalescence in a random background (Barton et al., 2004). This yields novel results. For example, in the strong selection case in which the selection strength *α* is proportional to the sample size *n* and both go to infinity (our scenario (iii)), the genealogy of the rare allele can be described in terms of a Cox-Ingersoll-Ross (CIR) diffusion with an initial Gamma distribution.

The concept of rare alleles in this paper and in Wakeley et al. (2023) is the same as the one considered by Joyce and Tavaré (1995) and Joyce (1995). It focuses on the counts of the alleles in a large sample rather than their relative frequencies in the population. In scenarios (ii) and (iii) we consider a fixed number *n*_1_ of the rare type 1 when the sample size *n* tends to infinity. Joyce and Tavaré (1995) considered rare alleles in a large sample drawn from the stationary distribution of a *d*-dimensional Wright-Fisher diffusion with selection and mutation. They showed that the counts of rare alleles, from different latent mutations in our terminology, have approximately independent Poisson distributions with parameters that do *not* depend on the selection parameters, and that the Ewens sampling formula describes their distribution. Their model with *d* = 2 and genic selection corresponds to our scenario (ii). Our results for very strong selection (*α* → ∞) in scenario (iii) differ from those of Joyce and Tavaré (1995) in that the rare-allele sampling probabilities (11a), (11b) and (11c) do depend on selection. Interestingly, the number of latent mutations given *n*_1_ still follows the Ewens sampling formula when lim_*n→∞*_ *α/n* ∈ (−∞, 1). But this is not true when lim_*n→∞*_ *α/n* ∈ (1, ∞), in which case the number of latent mutations is always *k*_1_ ≡ 1.

Some of our results for rare alleles have empirical relevance, specifically those for scenario (ii) including their robustness to time-varying population size demonstrated in Section 3.4, and those for scenario (iii) with 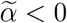. In scenario (ii), as *n* increases for fixed but arbitrary *α*, the distributions of latent mutations and the ages of those latent mutations become identical to those for neutral alleles described in Wakeley et al. (2023). Our results also show that selection does have an effect in this case, but it is only to raise or lower the rare-allele sampling probability (10) by the constant factor *C* for every value of *n*_1_. This relative insensitivity to selection suggests confidence in using rare alleles for demographic inference and genome-wide association studies (O’Connor et al., 2015; Nait Saada et al., 2020; Zaidi and Mathieson, 2020). Slatkin and Rannala (1997b), who obtained the Ewens sampling formula result for rare deleterious alleles by assuming they evolve independently according to a linear birth-death process, cf. Slatkin and Rannala (1997a), suggested that deviations from this neutral prediction at two human-disease-associated loci were due to population growth. Reich and Lander (2001) made a similar argument for a number of other disease-associated loci starting from the mutation-selection balance model of Hartl and Campbell (1982) and Sawyer (1983) which also gives the Ewens sampling formula result for rare disease alleles.

Our exploration of time-varying populations in Section 3.4, namely the robustness of the Ewens sampling formula result for the number of latent mutations, suggests that rare alleles may not always be well suited for demographic inference. With only a mild constraint on the trajectory of population sizes through time, increasing the sample size will eventually make the distribution of latent mutations of rare alleles look as if the population size has been constant at its current size. There is no doubt that demographic inferences improve as sample sizes increase. What Section 3.4 implies is that these improvements will not come from focusing exclusively on the lower end of sample allele frequencies (i.e. any fixed *n*_1_ as *n* → ∞). How relevant this is for a given sample will depend on the actual ages of its latent mutations and the degree of population-size change between those times and the present. To illustrate, consider the *O*(1*/n*) ages of latent mutations under the exponential growth model with rate *β*. If *β/n* ≪ 1, the ancestral process of tracing back to these mutations will be complete before the population has changed much in size and the results of Section 3.4 will hold. But this is clearly not the case for the *gnomAD* data in Wakeley et al. (2023) and Seplyarskiy et al. (2023). The distribution of *n*_1_ in the non-Finnish European sample with *n* = 114*K* is well fit by *β/n* = 3. See for example Fig. 3 in Seplyarskiy et al. (2023). Sample sizes would need to be orders of magnitude greater for the results in Section 3.4 to hold in this case.

Scenario (iii) with 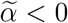 is applicable to strongly deleterious alleles. An appreciable fraction of new mutations are strongly deleterious (Eyre-Walker and Keightley, 2007; Kim et al., 2017; Weghorn et al., 2019; Dukler et al., 2022). Previous theoretical work includes Nei (1968), who found a gamma density analogous to ours in Lemma 1 but for the population allele frequency of partially recessive lethal mutations, and Charlesworth and Hill (2019), who used Nei’s approximation to derive the negative binomial distribution for *n*_1_, our (11a). In this case, (12a) shows that the sampling probabilities of rare alleles fall off quickly as *n*_1_ grows: each additional copy of *A*_1_ in the sample lowers its probability by a factor of 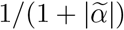 compared to the neutral case. Even so, the distribution of *k*_1_ given *n*_1_ follows the Ewens sampling formula. Hartl and Campbell (1982) and Sawyer (1983) obtained similar results by assuming that both selection and mutation are strong. Our analysis of scenario (iii) with 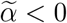 also shows that latent mutations of rare strongly deleterious alleles are especially young: selection speeds up the ancestral process of latent mutation by a factor of 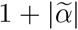 on top of the factor of *n* already present under neutrality. This is most easily seen by comparing the first two lines of (61) to the first two lines of (59).

Our results for scenario (i) with *α <* 0, which hold as *α* → −∞ for arbitrary sample size and alleles at any sample frequencies, are also applicable to strongly deleterious alleles. They are similar to the results just discussed for scenario (iii) with 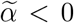. We expect that our results for very strong positive selection, i.e. scenario (i) with *α >* 0 and scenario (iii) with 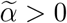, will be of limited applicability. Mutations to strongly positively selected alleles are uncommon and observing such an allele a small number of times in a very large sample would be exceedingly unlikely.

Many open questions remain. Joyce (1995) obtained a result similar to that of Joyce and Tavaré (1995), for a Wright-Fisher diffusion with selection and infinite-alleles mutation. This diffusion process is a particular case of the Fleming-Viot process (see Ethier and Kurtz (1993) for a review) and it has a unique stationary distribution denoted *ν*_selec_. Joyce (1995) considered a large sample of size *n* drawn from *ν*_selec_. Let 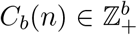 be the first *b* allele counts in a sample of size *n* drawn from the stationary distribution, and *K*_*n*_ be the total number of alleles in the sample. Joyce (1995) showed that for any fixed *b*, the distribution of (*C*_*b*_(*n*), *K*_*n*_) under *ν*_selec_ is arbitrarily close to that under the neutral model. It would be interesting to know if analogous results for our scenario (iii) also hold for the infinite allele model. In particular, is there a threshold for the selection strength relative to *n* that controls whether selection is washed out or not in the limit as *n* → ∞?

For time-varying populations, little is known in scenario (iii). For example, will the assumptions in Proposition 3 hold for a general demographic function? Will there be a phase transition for the value of *y*_*∗*_ in terms of 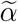 and if so, what will determine the phase transition? Also, both our results and those of Joyce and Tavaré (1995) and Joyce (1995) are for the infinite-population diffusion limit. Further consideration of the issues raised in Section 2.1.1 is needed to assess the relevance of these results to various kinds of finite populations.

The critical case 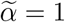 in scenario (iii) is omitted in this paper. Results for this case are expected to lie between those of 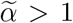 and 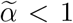, and require more in-depth asymptotic analysis. For example, one can first obtain asymptotic results for the hypergeometric function in (A.6a)-(A.6b) for the case 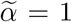, and then follow the argument in Lemma 1 to obtain the asymptotic of the expectation 𝔼_**n**_[*p*_0_] as *n*_2_ → ∞ in this critical case. Lemma 1 asserts that 𝔼_**n**_[*p*_0_] = *O*(1) when 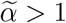 and 𝔼_**n**_[*p*_0_] = *O*(1*/n*_2_) when 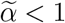. We conjecture that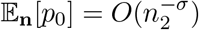 for some *σ* ∈ (0, 1) in the critical case.

Finally, we have ignored the possibility of spatial structure. Spatially heterogeneous populations in which reproduction rates, death rates, mutation rates and selection strength can depend both on spatial position and local population density present challenges. This is because the population dynamics now take place in high or infinite dimension (Hallatschek and Nelson, 2008; Barton et al., 2010; Durrett and Fan, 2016; Louvet and Véber, 2023; Etheridge et al., 2023). For example, the spatial version of (1), the stochastic Fisher-Kolmogorov-Petrovsky-Piscunov (FKPP) equation introduced by Shiga (1988), is a stochastic partial differential equation that arises as the scaling limit of various discrete models under weak selection (Müller and Tribe, 1995; Durrett and Fan, 2016; Fan, 2021). Under the stochastic FKPP, Hallatschek and Nelson (2008) and Durrett and Fan (2016) studied the backward-time lineage dynamics of a single sample individual, conditioned on knowing its type. It would be interesting to see if our results in this paper can be extended to spatial stochastic models with selection.

## Acknowledgements

We thank Alison Etheridge for raising the question about the applicability of our limiting results to Wright-Fisher reproduction (cf. Section 2.1.1). We also thank Shamil Sunyaev, Evan Koch and Joshua Schraiber for helpful discussions, and Daniel Rickert and Kejia Geng for assistance in producing the figures. Finally, we thank two anonymous reviewers for their insightful comments. This research was partially supported by National Science Foundation grants DMS-1855417 and DMS-2152103, and Office of Naval Research grant N00014-20-1-2411 to Wai-Tong (Louis) Fan.

## Appendix A. Asymptotic approximations used in the text

From the series expansion for a ratio of gamma functions with a common large parameter, 6.1.47 in Abramowitz and Stegun (1964) or equation (1) in Tricomi and Erdélyi (1951), we have

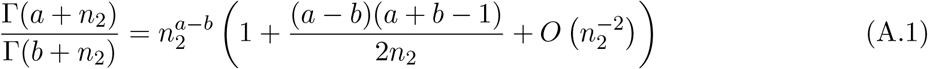

for constants *a* and *b* which will depend on the application. For example, we can apply (A.1) twice in the sampling probability (4) when *n*_2_ is large: once with *a* = *n*_1_ + 1 and *b* = 1 (in the binomial coefficient) and once with *a* = *θ*_2_ and *b* = *θ*_1_ + *θ*_2_ + *n*_1_.

The confluent hypergeometric function is commonly defined in terms of the integral

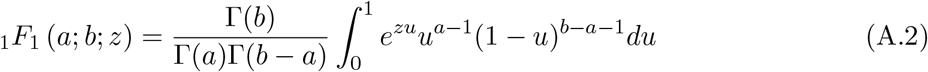

or in terms of the series

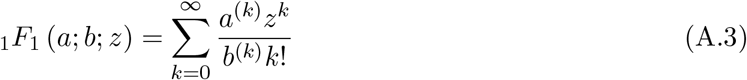

which converges for all *z* ∈ ℝ and *b > a >* 0, where *a*^(*k*)^ is the rising factorial *a*(*a* + 1) …(*a* + *k* − 1) with *a*^0^ = 1. Again *a* and *b* depend on the context, e.g. as in (3) and (4). The parameter *z* corresponds to the selection parameter *α*.

For large |*α*| and with constant *a* and *b*,

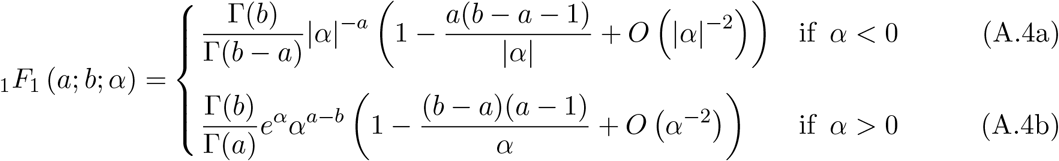

where the middle, neutral case is given only for completeness. Equation (A.4a) is from (4.1.2) in Slater (1960), and (A.4b) is from (4.1.6) in Slater (1960) or may be obtained from (A.4a) using Kummer’s first theorem which appears as (1.4.1) in Slater (1960).

For large *n*_2_, with constants *a, b* and *z*,

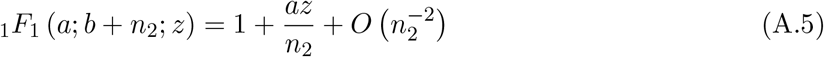

directly from (A.3).

For large *n*_2_ and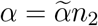, with constants *a* and *b*,

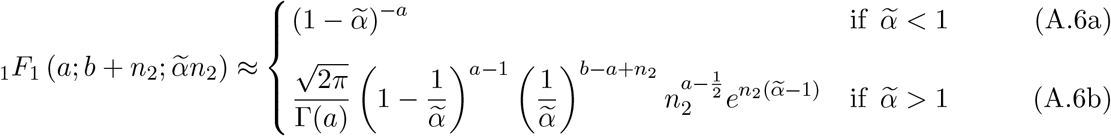

which we present only to leading order for simplicity. Equation (A.6a) follows from (A.3) and (A.6b) was obtained by applying Laplace’s method to the integral in (A.2) for this case.

## Appendix B. Proofs of Lemma 1, Lemma 1 and Lemma 1

*Proof of Lemma 1*. Part (ii) then follows from part (iii) with 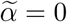, and the proof of part (i) follows from a similar argument.

To prove part (iii), we let *a* = *θ*_1_ + *n*_1_ for simplicity. The function 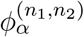 is a constant multiple of the function

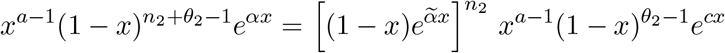

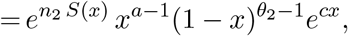

where the function *S* : [0, 1) → ℝ defined by 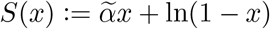

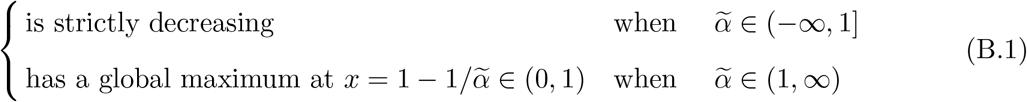

Part (iii) then follows from asymptotic expansion of integrals such as the Laplace method.

Let *x*^*∗*^ ∈ [0, 1] be the global maximum of the function *S*. Then *x*^*∗*^ = 0 when 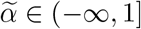 and 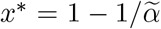 when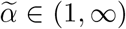. Fix an arbitrary *ϵ* ∈ (0, 1). There exists *δ* ∈ (0, 1) small enough such that 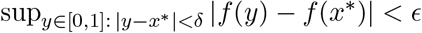 For each of the two cases, by (B.1), the ratio

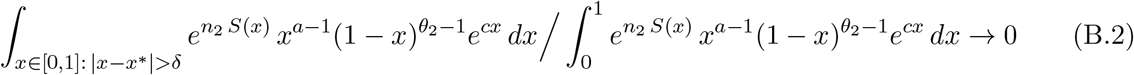

as *n*_2_ → ∞. For any *f* ∈ *C*_*b*_([0, 1]),

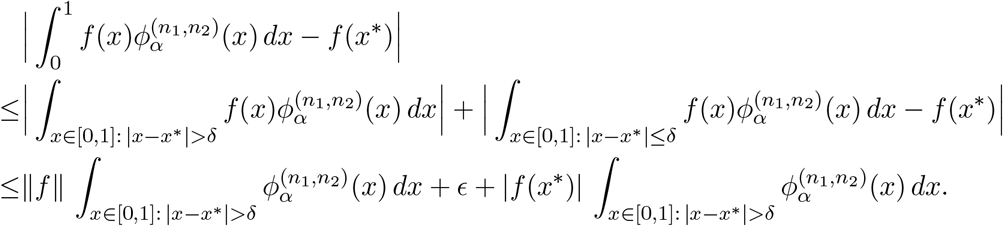

Hence by (B.2), lim 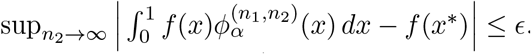. Since *ϵ >* 0 is arbitrary, we have shown that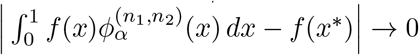 as *n*_2_ → ∞. □

*Proof of Lemma 1*. Convergence in distribution to a constant is equivalent to convergence in probability. Hence Lemma 1, except the last statement about the convergence in distribution of *n*_2_*p*_0_, follows from Lemma 1. As in the main text, *A* ≈ *B* below means *A/B* → 1 in the specified limit.

When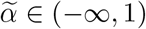, we let *a* = *n*_1_ + *θ*_1_ for simplicity. The probability density function of *np*_0_ under 𝕡_**n**_ is

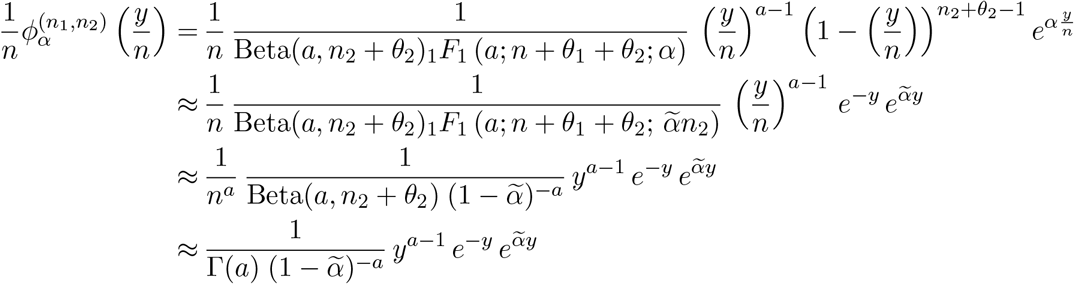

as *n*_2_ → ∞, where we used (A.6a) and then (A.1) in the last two approximations above. Hence the probability density function of *n*_2_*p*_0_ (under 𝕡_**n**_) converges pointwise to that of the 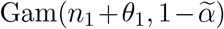 random variable. This implies the desired convergence in distribution. □

*Proof of Lemma 1*. Fix *t* ∈ ℝ_+_ and let *k* = [*N* (*N* − 1)*t/*2]. Suppose *A*(*k*) is the number of type 1 at step *k* of the discrete-time Moran process. Direct calculations from (13) and (14) show that, as *N* → ∞,

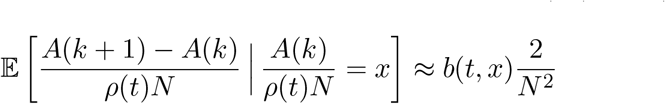

and

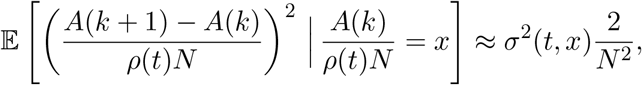

where 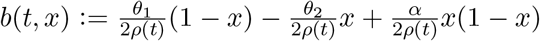 and 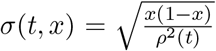are the coefficients in (44). The condition on *ρ* guarantees that the SDE (44) has a unique weak solution and that the desired weak convergence follows from standard (martingale problem) method; for reference see Stroock and Varadhan (1979, Chapter 11). □

## Appendix C. Events in the conditional ancestral selection graph

Here we show how the minimal conditional ancestral process in Section 4 is obtained from the full conditional ancestral process. To begin, we assume that at some time in the conditional ancestral process there were *r*_1_, *r*_2_, *v*_1_ and *v*_2_ real and virtual lineages of type 1 and type 2. The associated sampling probability is *q*_*o*_(*r*_1_, *r*_2_, *v*_1_, *v*_2_), the straightforward extension of (53) or (54) to include both type-1 and type-2 virtual lineages. How branching events are resolved depends on which allele is favored by selection. We begin here by assuming that *A*_2_ is favored, or *α <* 0. Grouping events by the types of lineages involved (real or virtual of type 1 or type 2) then by whether it is mutation, branching or coalescence gives fourteen possibilities which occur at the following rates.

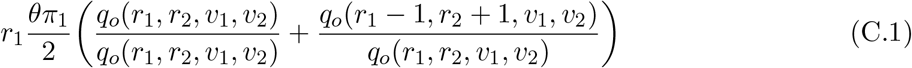

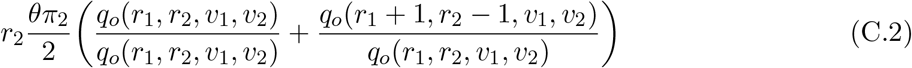

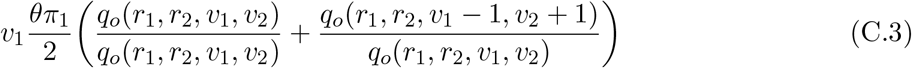

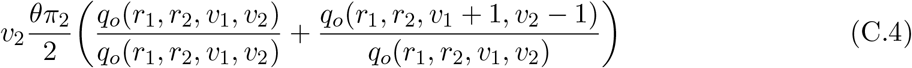

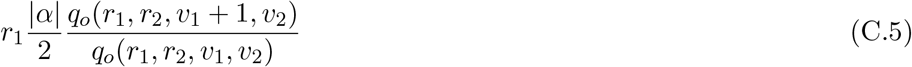

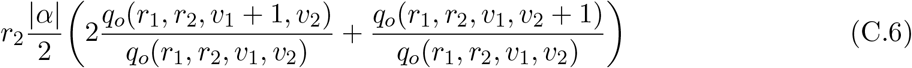

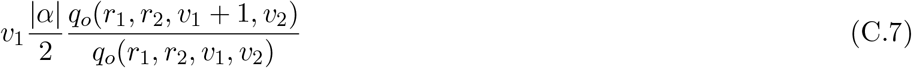

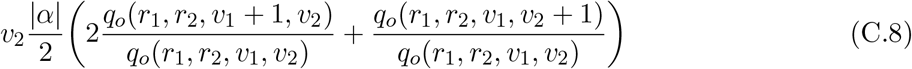

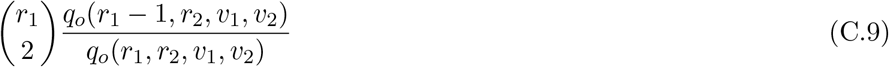

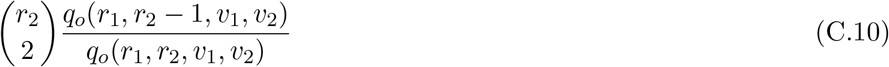

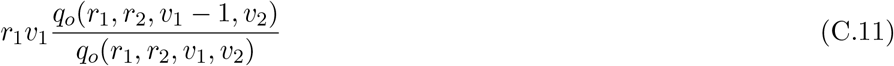

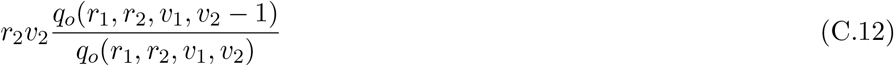

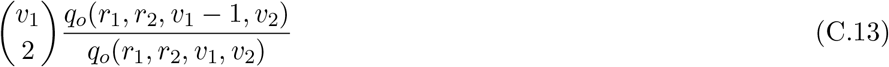

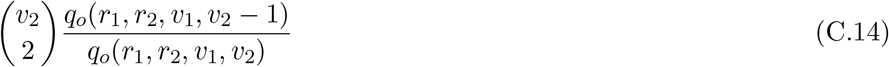

The sum of (C.1) through (C.14) is equal to the total rate of events in the unconditional ancestral process, (*r*_1_ + *r*_2_ + *v*_1_ + *v*_2_)(*θ* + |*α*| + *r*_1_ + *r*_2_ + *v*_1_ + *v*_2_ − 1)*/*2. Twenty-two distinct events 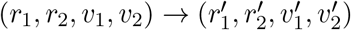 are represented, one for each of the ratios of sampling probabilities,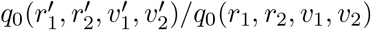. Note that the assumption of parent-independent mutation leads to the four kinds of spurious or empty mutation events in (C.1) through (C.4) which do not change the ancestral state of the sample 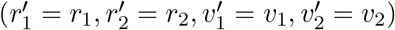. Also, only those events which have have non-zero probabilities of giving the data appear in (C.1) through (C.14); coalescent events between lineages with different types and type-*i* mutation events on type 3 − *i* lineages would make the data impossible.

Recall that the resolution of branching events depends on which allele is favored by selection. The events and their probabilities in (C.5) through (C.8) are just for the case *α <* 0, where *A*_2_ is the favored allele. Each branching event creates an incoming lineage and a continuing lineage, both of which may be of type 1 or type 2. Let (*I, C*) be the types of these lineages. In (C.5) and (C.7), only one of the four (*I, C*) pairs has non-zero probability of producing the data: (*I* = 1, *C* = 1) corresponding to the event (*r*_1_, *r*_2_, *v*_1_, *v*_2_) → (*r*_1_, *r*_2_, *v*_1_ + 1, *v*_2_). In (C.6) and (C.8), the possibility (*I* = 1, *C* = 1) is discarded as it would then be impossible for the descendant lineage to be of type 2. The other three possibilities have non-zero chances of producing the data, and associated events

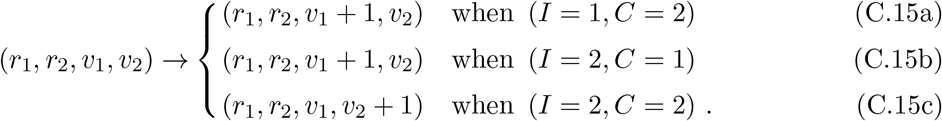

In contrast, if *α >* 0 then branching events on type-2 lineages are the ones for which only one of the four (*I, C*) pairs has non-zero probability of producing the data: (*I* = 2, *C* = 2) corresponding to the event (*r*_1_, *r*_2_, *v*_1_, *v*_2_) → (*r*_1_, *r*_2_, *v*_1_, *v*_2_ + 1). When *α >* 0, if the branching event occurs on a type-1 lineage, then in place of (C.15a), (C.15b) and (C.15c) we have

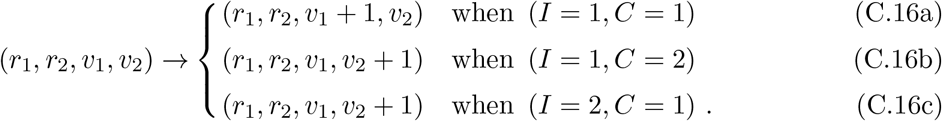

Therefore, when *α >* 0, (C.5) through (C.8) must be replaced with

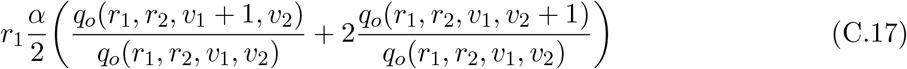

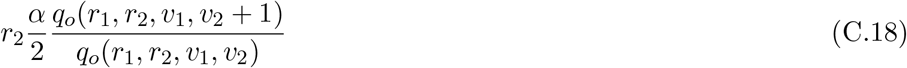

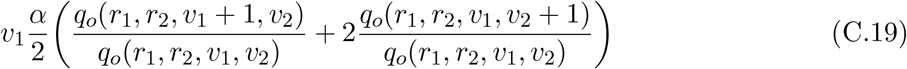

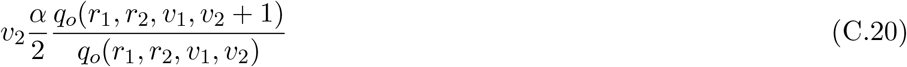

Equations (C.1) through (C.7) and (C.9) through (C.14) are the same for *α >* 0 and *α <* 0.

The simplifications discovered by Slade (2000a) and Fearnhead (2002) follow from the simple fact that each sampled lineage is either of type 1 or type 2. Slade (2000a) noticed that when both the descendant lineage and the incoming lineage have the favored type, the type of the continuing lineage does not matter so there is no need to introduce a new virtual lineage. Instead, these two possibilities can be collapsed into a single null event which does not change the numbers and types of ancestral lineages. That is, we can use

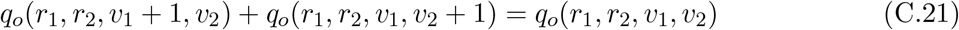

in (C.6), (C.8), (C.17) and (C.19). As a result, no type-2 virtual lineages will be created.

Along the same lines, Fearnhead (2002) noticed that when mutation is parent-independent there is no need to follow ancestral lineages once they have mutated, because the ancestral lineage could be of either type. Any such lineage can be removed from the ancestral process. Here we use

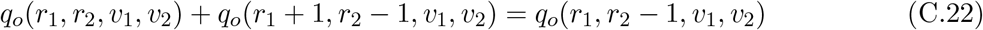

in (C.2), and other appropriate identities in (C.3) and (C.4). But we do not make use of this simplification in (C.1) because our specific goal is to model latent mutations in the ancestry of *A*_1_. These are actual mutations, where the ancestral type was *A*_2_. The remaining *A*_1_ → *A*_1_ empty mutations are null events, which do not change the numbers and types of ancestral lineages.

The conditional ancestral processes for *α <* 0 and *α >* 0 given by (55) and (56) in the main text each include just five kinds of (non-null) events. We obtain these by applying the simplifications of Slade (2000a) and Fearnhead (2002) then grouping events by their outcomes. For example, the coalescent events in (C.10) have effect *r*_2_ → *r*_2_ − 1, as do the combined mutations in (C.2) once the simplification of Fearnhead (2002) is applied. So these appear together as one kind of event, the fifth case in both (55) and (56).

We do not include null events in (55) and (56) since these by definition have no effect on the ancestral lineages. In the case *α <* 0, the null events are empty mutations on type-1 real lineages and branching events on type-2 real lineages where the incoming line is also of type 2. These occur with total rate *r*_1_*θπ*_1_*/*2 + *r*_2_|*α*|*/*2. In the case *α >* 0, the null events are empty mutations on type-1 real lineages and branching events on type-1 real lineages where the incoming line is also of type 1. These occur with total rate *r*_1_*θπ*_1_*/*2 + *r*_1_*α/*2.

